# The genome and lifestage-specific transcriptomes of a plant-parasitic nematode and its host reveal susceptibility genes involved in trans-kingdom synthesis of vitamin B5

**DOI:** 10.1101/2021.10.01.462558

**Authors:** Shahid Siddique, Zoran S. Radakovic, Clarissa Hiltl, Clement Pellegrin, Thomas J. Baum, Helen Beasley, Oliver Chitambo, Divykriti Chopra, Etienne G.J. Danchin, Eric Grenier, Samer S. Habash, M. Shamim Hasan, Johannes Helder, Tarek Hewezi, Julia Holbein, Martijn Holterman, Sławomir Janakowski, Georgios D. Koutsovoulos, Olaf P. Kranse, Jose L. Lozano-Torres, Tom R. Maier, Rick E. Masonbrink, Badou Mendy, Esther Riemer, Mirosław Sobczak, Unnati Sonawala, Mark G. Sterken, Peter Thorpe, Joris J.M. van Steenbrugge, Nageena Zahid, Florian Grundler, Sebastian Eves-van den Akker

## Abstract

Plant-parasitic nematodes are a major, and in some cases a dominant, threat to crop production in all agricultural systems. The relative scarcity of classical resistance genes highlights a pressing need to identify new ways to develop nematode-resistant germplasm. Here, we sequence and assemble a high-quality genome of the model cyst nematode *Heterodera schachtii* to provide a platform for the first system-wide dual analysis of host and parasite gene expression over time, covering all major stages of the interaction. This novel approach enabled the analysis of the hologenome of the infection site, to identify metabolic pathways that were incomplete in the parasite but complemented by the host. Using a combination of bioinformatic, genetic, and biochemical approaches, we show that the highly atypical completion of vitamin B5 biosynthesis by the parasitic animal, putatively enabled by a horizontal gene transfer from a bacterium, is critically important for parasitism. Knockout of either the plant-encoded or the now nematode-encoded steps in the pathway blocks parasitism. Our experiments establish a reference for cyst nematodes, use this platform to further our fundamental understanding of the evolution of plant-parasitism by nematodes, and show that understanding congruent differential expression of metabolic pathways represents a new way to find nematode susceptibility genes, and thereby, targets for future genome editing-mediated generation of nematode-resistant crops.

## Introduction

Plant-parasitic nematodes are a major threat to crop production systems all around the world, and in some crops (e.g. soybean) are the dominant pathogen of any kind (Savary et al. 2019; Abd-Elgawad, Askary, and Others 2015). They are estimated to cause losses of up to 25% each year, with a value of more than 80 billion US dollars (Nicol et al. 2011). The major contributors to these losses are the notoriously difficult to control obligate biotrophic sedentary endo-parasites: a remarkable group of parasites that live inside, and feed on living host root cells.

Cyst nematodes, one of the two major groups of sedentary endo-parasites, are a perennial constraint on crop production. Their control is difficult due to their complex biology. A cyst, formed from the body wall of a dead female, contains hundreds of eggs that can remain dormant in the soil for many years. Eggs contain a fully-formed, but sexually undetermined, second stage juvenile (J2) that will hatch in response to various host-derived cues (Baunacke 1922). Hatching is a critical decision: the J2 will migrate through the soil to find a host, invade the root through a combination of mechanical disruption using a needle-like stylet and enzymatic secretions, and move towards the vascular cylinder – all without feeding. Once J2s reach the vascular cylinder, their behaviour changes (stylet thrusts become more “exploratory”), and a single cell is “chosen” for feeding site development (Wyss 1992). To manipulate plant-development and immunity, the nematode injects a cocktail of proteins and other molecules, termed “effectors’’ into the plant cell cytoplasm (Lilley et al. 2018) and surrounding apoplast (Eves-van den Akker et al. 2014). A majority of these effectors originate from two sets of gland cells (one dorsal and two subventral) and cause an existing plant cell, often procambial, to re-differentiate into a syncytial feeding site that is unlike any other tissue in the plant (Szakasits et al. 2009).

Host cell manipulation is rapid and profound: within just a few days, the cell cycle arrests at G2 phase, the vacuole reduces in size and fragments, the nucleus greatly enlarges, the cytoplasm becomes enriched in organelles by extensive proliferation of the rough and smooth endoplasmic reticulum, ribosomes, mitochondria and plastids (chloroplasts and amyloplasts). The cell wall is locally degraded to promote protoplast fusion with hundreds of adjacent cells in an iterative manner (Golinowski, Grundler, and Sobczak 1996; Sobczak, Golinowski, and Grundler 1997; F. M. W. Grundler, Sobczak, and Golinowski 1998). Establishment of the feeding site is a point of no return: concurrent with establishing the feeding site, cyst nematodes become sedentary. If at any time during the interaction the syncytium dies, so does the nematode (Raski and Others 1950).

Unusually for an animal, cyst nematode sex is determined during parasitism. It is understood that the sexually undetermined juvenile with access to ample nutrition (from a well-formed feeding site) will preferentially develop into a female, while those with limited nutrition preferentially develop into males, stop feeding, and regain motility and search for a mate (F. Grundler, Betka, and Wyss 1991; Anjam et al. 2020). The obligate sexual cyst nematode females will ultimately fill with eggs, their body wall will tan and harden, and the resultant cyst will drop off the root, and their eggs can remain dormant in the soil for several years.

Cyst nematodes (particularly the *Heterodera* genus) are notoriously difficult to control due, at least in part, to the relative scarcity of commercially viable resistance sources. Resistance can be achieved by either gain-of-function “resistance genes” (plant genes that block the parasite/feeding site development in some way), or loss-of-function of “susceptibility genes” (plant genes that are necessary for feeding site development/nematode infection). Given the relative scarcity of classical resistance genes against Heterodera (Yan and Baidoo 2018; Kumar et al. 2021) there is a pressing need to better understand the biology of host infection and identify nematode susceptibility genes. The beet cyst nematode, *Heterodera schachtii*, is the obvious choice in which to build this understanding because: a) it can infect the model plant *Arabidopsis thaliana*, b) it is an important agricultural pest on sugar beet and brassicaceous crops in its own right, and c) its close relative the soybean cyst nematode, *H. glycines*, is one of the severely economically damaging plant-parasitic nematodes worldwide.

Here we sequence and assemble a high-quality genome of the beet cyst nematode *H. schachtii* to provide a platform for the first system-wide and simultaneous analysis of host and parasite gene expression across time, covering all major stages of the interaction. To focus on novel aspects of biotrophy that may reveal nematode susceptibility genes, we analysed metabolic pathways that were incomplete in the parasite, but complemented by the host. Using a combination of bioinformatic, genetic, and biochemical approaches, we highlight the highly atypical completion of the vitamin B5 (pantothenate) biosynthesis pathway by the nematode parasite that was most likely enabled by a horizontal gene transfer from a bacterium. Importantly, knockout of either the plant-encoded or the now nematode-encoded steps in the pathway blocks parasitism. Our experiments establish a reference for cyst nematodes, use this as a platform to further our fundamental understanding of the evolution of plant-parasitism, and show that the understanding of congruent differential expression of metabolic pathways represents a new way to find nematode susceptibility genes, and thereby targets for the generation of genome edited crops resistant to nematodes.

## Results and discussion

### Sequencing and assembly of the *H. schachtii* genome

We measured (Figure S1), sequenced (BioProject PRJNA722882), and assembled the genome of *H. schachtii* (population Bonn) using a combination of flow cytometry, Pacific Biosciences sequencing, and Illumina sequencing. *H. schachtii* has the largest genome (160-170 Mb) of any cyst nematode measured/sequenced to date (Table S1). It was sequenced to 192-fold coverage using Pacific Biosciences sequencing (fragment n50 of 16 kb), and 144-fold coverage using Illumina sequencing (150 bp Paired end reads). The final, polished, contamination-free (Figure S2), assembly (v1.2) included approximately 179 Mbp contained within 395 scaffolds: 90% of the sequence is contained on scaffolds longer than 281,463 bp (n=154). The assembly is a largely complete haploid representation of the diploid genome, as evidenced by core eukaryotic genes being largely present, complete and single copy (CEGMA 93.15% complete with an average of 1.12 copies each, and BUSCO (Eukaryota odb9) 79% complete with 8.2% duplicated).

### The trans-kingdom, life stage-specific, transcriptomes of *H. schachtii* and *A. thaliana* provide a holistic view of parasitism

We devised a sampling procedure to cover all major life stages/transitions of the parasitic life cycle to generate a simultaneous, chronological, and comprehensive picture of nematode gene expression, and infection-site-specific plant gene expression patterns. We sampled cysts and pre-infective second-stage juveniles (J2s), as well as infected segments of *A. thaliana* root and uninfected adjacent control segments of root at 10 hours post infection (hpi - migratory, pre-establishment of the feeding site), 48hpi (post establishment of the feeding site), 12 days post infection females (dpi - virgin), 12 dpi males (differentiated but pre-emergence), and 24 dpi females (post mating), each in biological triplicate (Figure 1A). We generated approximately 9 billion pairs of 150 bp strand-specific RNAseq reads (Table S2) covering each stage in biological triplicate (for the parasite and the host): in the early stages of infection we generated over 400 million reads per replicate, to provide sufficient coverage of each kingdom.

**Figure 1:**
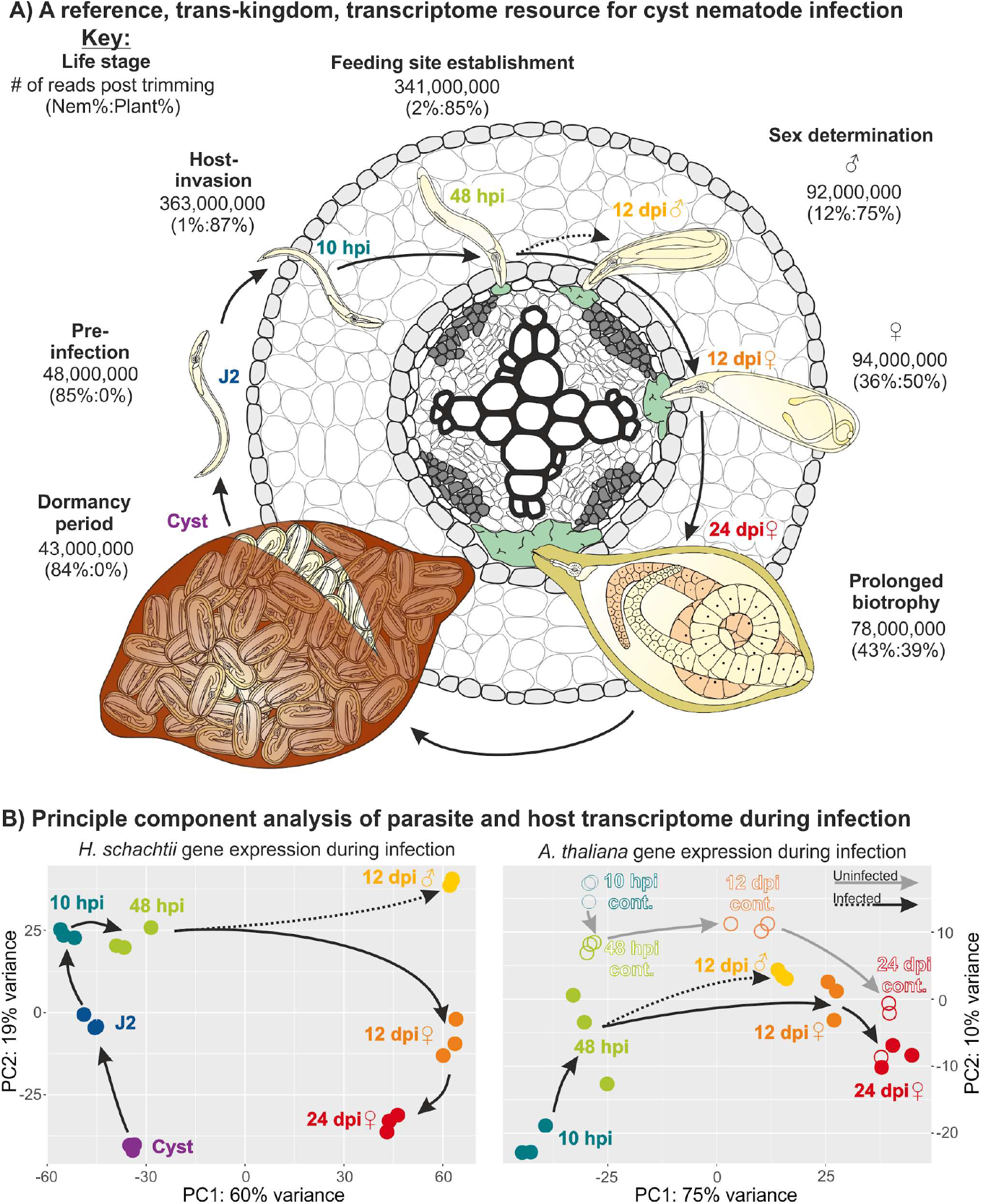
Trans-kingdom, life stage-specific, transcriptome of *H. schachtii* and *A. thaliana*. **A**) Schematic representation of the life cycle of *H. schachtii* infecting *A. thaliana*, highlighting the 7 stages sampled in this study. For each stage, the average number of trimmed RNAseq read pairs per replicate is shown, with the proportion of reads mapping to either parasite or host in parentheses. **B**) Principle components 1 and 2 for *H. schachtii* and *A. thaliana* expression data are plotted. Arrows indicate progression through the life cycle/real time. Hours post infection (hpi), Days post infection (dpi).

Strand-specific RNAseq reads originating from host and parasite were deconvoluted by mapping to their respective genome assemblies (*H. schachtii* v.1.2 and TAIR10). For the parasite, approximately 500 million Illumina RNAseq read pairs uniquely mapping to the *H. schachtii* genome were used to generate a set of 26,739 gene annotations (32,624 transcripts - detailed further in the next section), approximately 77% of which have good evidence of transcription in at least one life stage (≥10 reads in at least one rep). Similarly for the host, approximately 2.8 billion Illumina RNAseq read pairs uniquely mapping to the *A. thaliana* genome show that approximately 77% of the 32,548 gene models have good evidence of transcription in at least one stage (≥10 reads in at least one rep, even though we only sampled roots). A principal component analysis of the host and parasite gene expression data offers several insights into the parasitic process. Principle component 1 (60% of the variance) and 2 (19% of the variance) of the parasite recapitulate the life cycle in PCA space (Figure 1B). The 12 dpi female transcriptome is more similar to the 24 dpi female transcriptome than to the 12 dpi male transcriptome. Principle components 1 (75% of the variance) and 2 (10% of the variance) of the host show that the greatest difference between infected and uninfected plant tissue is at the early time points (10 hpi), and that the transcriptomes of infected and uninfected plant material converge over time. A 12 dpi male syncytium transcriptome is roughly intermediate between a control root transcriptome and a 12 dpi female syncytium transcriptome. By comparing both principal component analyses, we can see that a relatively small difference in the transcriptomes of the feeding sites of males and females is amplified to a relatively large difference in the transcriptomes of the males and females themselves (Figure 1B).

### The consequences, and possible causes, of large-scale segmental duplication in the Heterodera lineage

To understand the evolutionary origin(s) of the relatively large number of genes in *H. schachtii* in particular, and *Heterodera* spp. in general, we analysed the abundance and categories of gene duplication in the predicted exome. Compared to a related cyst nematode, *Globodera pallida* (derived using comparable methodology and of comparable contiguity) the exomes of *H. schachtii* and *H. glycines* are characterised by a relatively smaller proportion of “single copy” genes (as classified by MCSanX toolkit (Wang et al. 2012), and a relatively greater proportion of “segmental duplications” (at least 5 co-linear genes with no more than 25 genes between them), with relatively similar proportions of “dispersed duplications” (two similar genes with more than 20 other genes between them), “proximal duplications” (two similar genes with less than 20 other genes between them), and “tandem duplications” (two similar genes that are adjacent) (Figure 2A). Genes classified as segmentally duplicated are clustered into “islands” (Figure 2B) of varying gene duplication depth from 2 to 131. Setting an arbitrary duplication depth threshold of 10, approximately half of all segmentally duplicated genes (2,273/4,547) are grouped into 58 “islands” across the genome.

**Figure 2.**
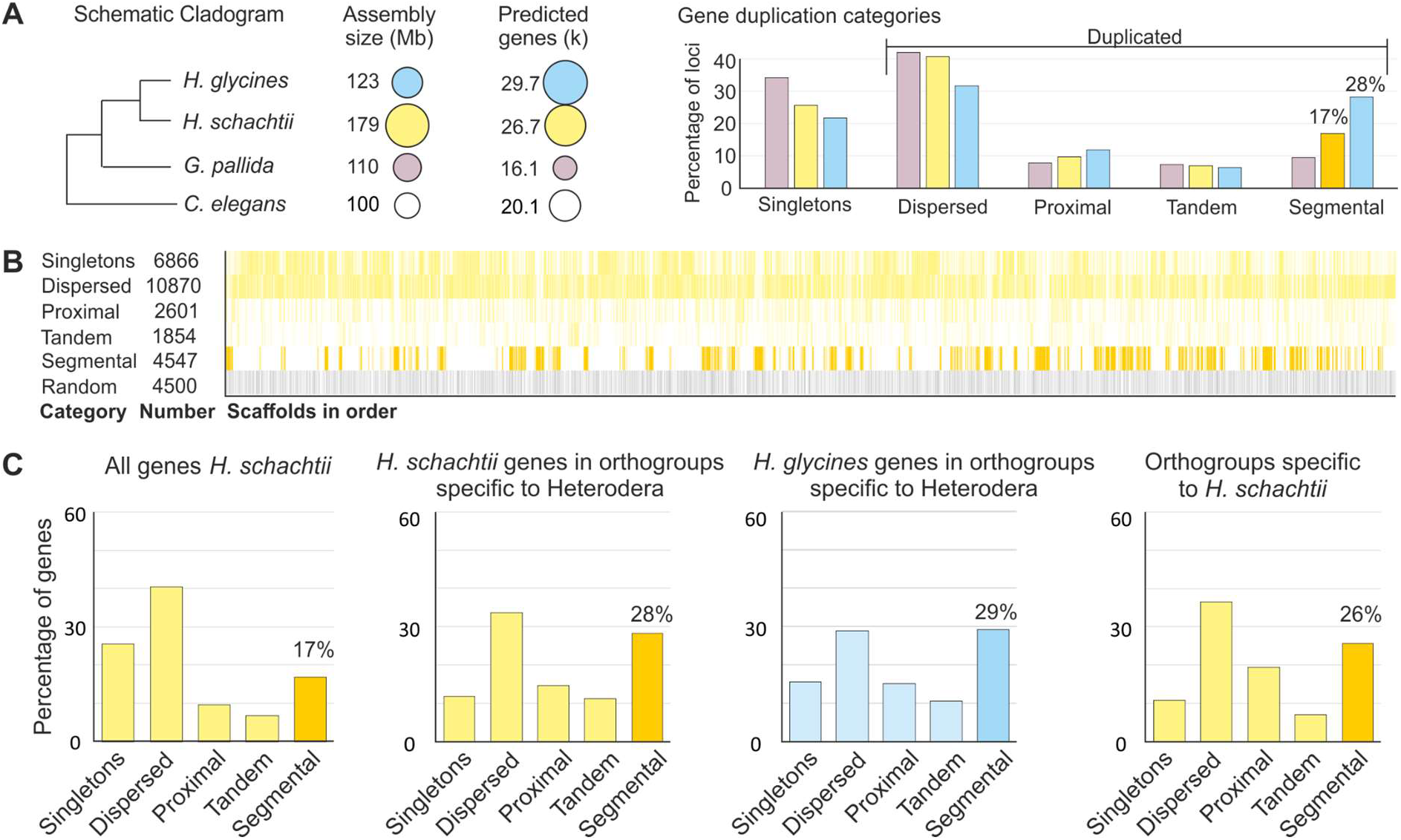
Large-scale segmental duplications in the Heterodera lineage acted on old new genetic capital. **A**) Schematic cladogram, assembly size, number of predicted genes, and gene duplication categories are shown for *H. schachtii* (yellow), *H. glycines* (blue), *G. pallida* (purple), and *C. elegans* (white). **B**) The genomic distribution of gene duplication categories. Each bar represents one gene. **C**) Gene duplication plots, from left to right: all *H. schachtii* genes, *H. schachtii* genes in orthologous gene clusters categories specific to Heterodera, *H. glycines* genes in orthologous gene clusters categories specific to Heterodera, and *H. schachtii* genes in orthologous gene clusters categories specific to *H. schachtii*.

To understand when this large-scale segmental duplication occurred, and what genetic capital it has acted on, we cross-referenced the gene duplication analyses with a recent orthologous gene clustering of 61 species across the phylum *Nematoda* (Grynberg et al. 2020). Interestingly, most (434/833) of the orthologous gene clusters that contain *H. schachtii* genes classified as segmentally duplicated also contain *H. glycines* genes classified as segmentally duplicated. We therefore conclude many are likely bona fide, and that at least half of the modern day segmentally duplicated regions of the *H. schachtii* genome were present in the last common ancestor with *H. glycines*.

Interestingly, gene families that have arisen in the *Heterodera* lineage (i.e. contain more than one *H. schachtii* or *H. glycines* gene, with no representatives from all 61 other proteomes in the analysis) are numerous (1,877 orthogroups, accounting for 2,072 genes in *H. schachtii*) and are also enriched in segmental duplications (28% vs 17% for *H. schachtii*; Figure 2C). Finally, *H. schachtii* specific gene families (i.e. those that only contain more than one *H. schachtii* gene, with no representatives from all 61 other proteomes in the analysis, including *H. glycines*) are also preferentially located in regions of the genome that have been segmentally duplicated (26% vs 17% of all genes; Figure 2C). We reason that some of these segmental duplications must have arisen relatively recently (i.e. since divergence from *H. glycines*) because although it is not certain whether *H. schachtii*-specific genes arose in the *H. schachtii* lineage or were lost in the *H. glycines* lineage, if these gene families were segmentally duplicated before the divergence of *H. schachtii* and *H. glycines*, then *H. glycines* would need to have independently lost both segments to not be in the orthogroup, and this seems unlikely for all cases. Taken together, we conclude that substantial segmental duplication started in the *Heterodera* lineage after the split from *Globodera*, was still active in the *H. schachtii* lineage after the split from *H. glycines*, and has acted on old and new genetic capital.

To understand the consequences, and possible causes, of these large scale segmental duplications, we adopted two complementary approaches: i) a targeted approach to examine what impact this may have had on genes known to be involved in parasitism, and ii) a non-targeted approach to examine Gene Ontology (GO) terms that are enriched in segmental duplications and/or gene families that are segmentally duplicated. We therefore additionally annotated the predicted proteome of *H. schachtii* to identify putatively secreted proteins (2,669 genes, 3,138 transcripts - Table S3), putative orthologues of known effectors (38 families, 209 genes, 245 transcripts - Table S6), genes putatively acquired by horizontal gene transfer (HGTs, 263 genes, 319 transcripts - Table S7), putative cell wall degrading enzymes (CWDEs, 9 families, 39 genes, 45 transcripts - Table S6), and PFAM domains (to assign Gene Ontology terms, 11,836 genes, 14,705 transcripts - Table S3).

In the targeted approach, we find preferential duplication of proper subsets of the secretome and the HGTs that are directly implicated at the plant-nematode interface (i.e. effectors and CWDEs, respectively; Figure 3A). For example, secreted proteins are not generally more duplicated than the rest of the genes in the genome, but effector genes are - including a substantial proportion in segmental duplications (approx. 1/5th). Similarly, putative HGT events are not generally more duplicated than the rest of the genes in the genome, but the cell wall degrading enzymes are (zero single copy genes; Figure 3A). In the non-targeted approach, we find that genes in segmental duplications are enriched in a very small set of highly-related GO terms involved in nucleic acid synthesis/manipulation and proteolysis (Figure 3B). Many of the PFAM domains underlying these enriched annotations are associated with DNA replication/integration of viruses/transposons and, remarkably, they tend to be located towards one or both edges of “islands” of segmental duplication (Figure 3C). The highly statistically significant functional enrichment of GO terms involved in DNA integration (adj. p-value = 7.46E^-92^), DNA biosynthetic process (adj. p-value = 1.53E^-11^), viral DNA genome packing (adj. p-value = 3.86E^-20^), and DNA replication (adj. p-value = 4.67E^-06^), coupled with their tendency to be positioned at the edge of segmental duplications, leads us to hypothesise that these functions may have played a role in the large-scale segmental duplication in this lineage.

**Figure 3.**
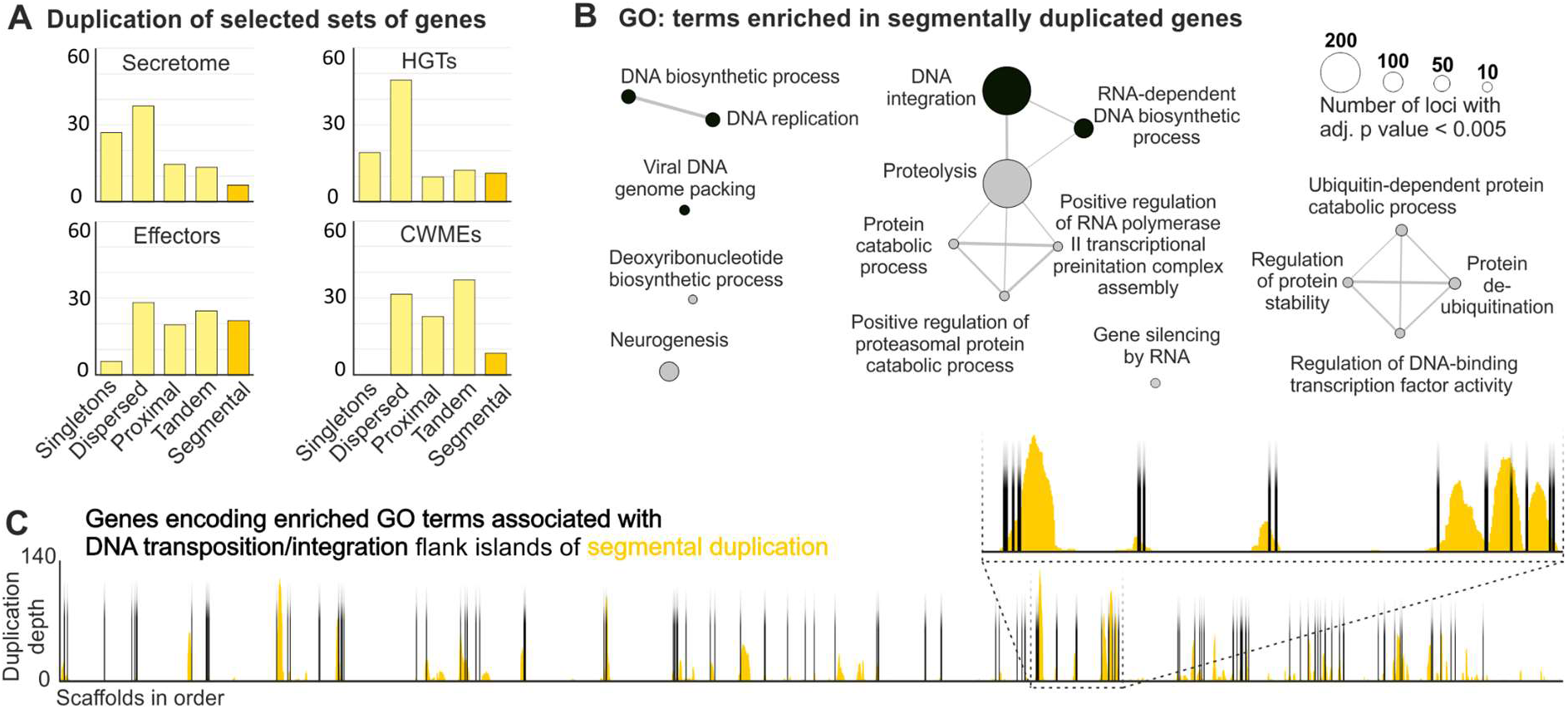
Islands of segmentally duplicated genes are flanked by genes predicted to be involved in DNA integration/transposition. **A**) Gene duplication plots for all putatively secreted proteins (secretome), effectors, genes putatively acquired via Horizontal Gene Transfer (HGTs), and genes encoding Cell Wall Modifying Enzymes (CWMEs). **B**) GO enrichment of segmentally duplicated genes in *H. schachtii*. **C**) Orange bars indicate duplication depth of segmentally duplicated genes across the genome. Black bars indicate the position of genes encoding enriched GO terms associated with DNA transposition/integration in segmental duplications.

### Differential expression analysis reveals contrasting evolutionary histories of host and parasite genes deployed at specific times

To understand how host and parasite genes are expressed during the parasitic life cycle, only those RNA-seq reads that are uniquely mapped to the host genome, not the parasite genome, and vice versa, were used to identify host and parasite genes that were significantly differentially expressed between conditions (Log2FC > +0.5 or < −0.5 and adj. p-value <0.01, Tables S4 and S5). Under these criteria, 69% (18,536) of the nematode genes, and 59% (19,071) of *A. thaliana* gene, are differentially regulated during the course of infection. To focus on the local infection specific host response (i.e. removing changes associated with time/systemic changes), we selected only the subset of the *A. thaliana* transcriptome, 25% (8,351 genes), that is differentially expressed between infected and uninfected tissues at each time point (hereafter referred to as differentially expressed genes). A majority of host and parasite differentially expressed genes (81% and 94% respectively) were assigned to super clusters that describe different stages, or groups of stages, of the infection cycle (29 and 28 super clusters for host and parasite, respectively - e.g. Figure S3, and Tables S3 and S4). For the host, the three largest clusters are of comparable size (10hpi_48hpi, 12dpi fem._24dpi, and 10hpi_48hpi_12dpi male), and together account for over 2/5ths of all differentially expressed genes. For the parasite, the cluster with the most differentially expressed genes has a peak at 12 dpi males. It contains nearly 1/5th of all differentially expressed genes, and is over twice as large as the next largest cluster (cyst). This is noteworthy because males and cysts have generally been overlooked in previous differential expression analyses in the literature, and yet these two life stages alone account for nearly 1/3rd of all differentially expressed genes. Interestingly, genes in the male-specific cluster are non-randomly distributed across the genome (for example, >31% of genes on scaffold 23 (115/364) are in the male specific cluster, hypergeometric test, adj. p-value = 1.51E^-25^ (Table S5)). Male-specific genes are often located in segmentally duplicated “islands”, even when the scaffold as a whole is not enriched, because 30% of all male-specific genes have sequence similarity to ADI82807.1 (860/2881), and 72% of genes with sequence similarity to ADI82807.1 are in segmental duplications (2235/3094). This gene family was noted as large (474), and male-expressed, in *G. pallida* (Cotton et al. 2014).

To analyse the evolutionary histories of clusters of genes that peak at specific life stages, we cross referenced these data with existing analyses of orthologous gene clusters of plants and nematodes. We focused on a subset of eight plant species (*Amborella trichopoda* (outgroup), the monocots *Hordeum vulgare* and *Zea mays*, the solanaceous *Solanum lycopersicum* and *S. tuberosum*, the fabaceous *Glycine max*, and the brassicaceous *Brassica rapa* (ssp. *rapa*) and *A. thaliana* - GreenPhylDB) and a subset of eight nematodes species (the free living nematode *Caenorhabditis elegans* (outgroup), the pine wilt nematode *Bursaphelenchus xylophilus*, the root-knot nematodes *Meloidogyne graminicola* and *M. hapla*, the potato cyst nematodes *G. pallida* and *G. rostochiensis*, and *H. glycines* and *H. schachtii* (Grynberg et al. 2020)). Six categories of orthologous clusters were considered for the parasite (nematodes, plant-parasites, root-knot nematodes and cyst nematodes, cyst nematodes, *Heterodera*, and *H. schachtii*), and six categories of orthologous clusters were considered for the plant (Magnoliopsida, Mesangiospermae (monocots and dicots), Pentapetalae, Rosids, Brassicaceae, and *A. thaliana*) (Figure S4).

We correlated the putative annotation, orthologue definition, and transcriptional clustering data to explore relatedness and functions of subsets of differentially expressed genes. Initially, we analysed the six differential expression clusters that describe the transition to biotrophy (J2, J2_10hpi, 10hpi, 10_48hpi, 48hpi, and J2_10_48hpi; Figure 4A). Of these six, five are significantly enriched in genes encoding putatively secreted proteins (hypergeometric test, false discovery rate (FDR) adj. p-value < 0.01), the exception being the J2 specific cluster, which also contains zero known effectors. This does not mean that no effectors are expressed in the J2 stage, but rather that none are specific: all effectors expressed in the J2 stage are also expressed at some other life stage such that they are clustered elsewhere.

**Figure 4.**
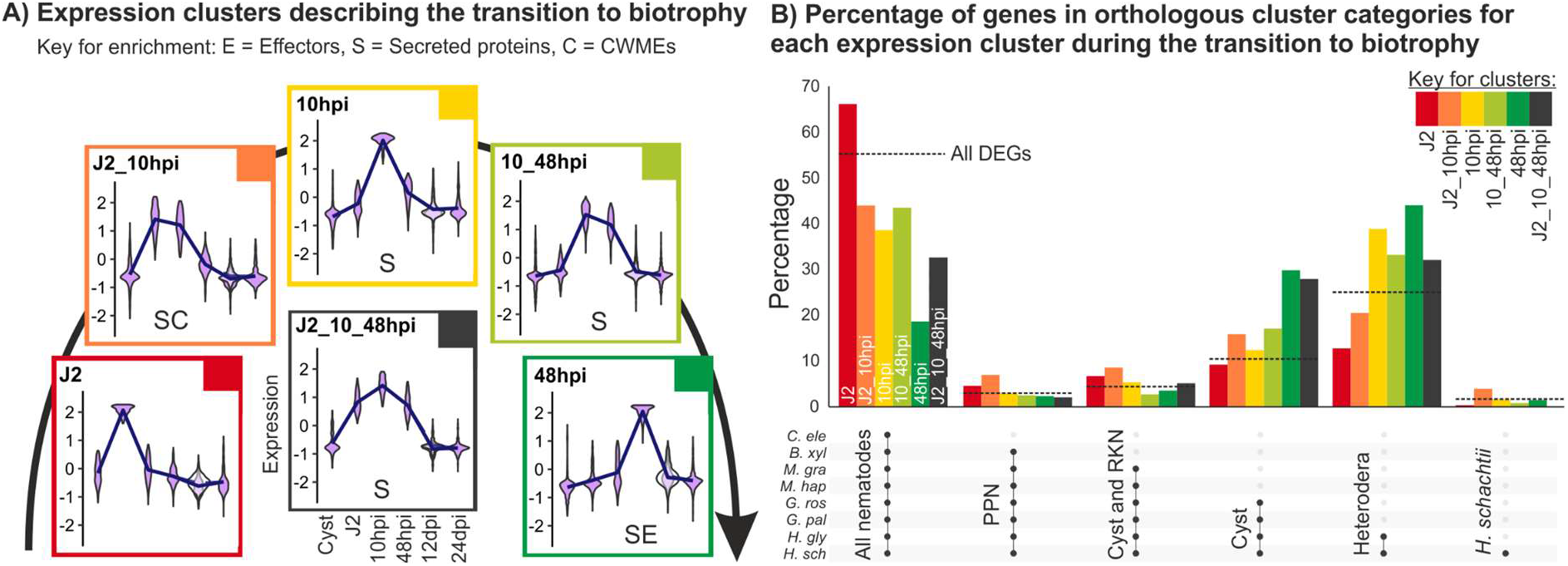
*H. schachtii* genes deployed during the transition to biotrophy are variously conserved with cyst nematodes. **A**) The expression profiles of six super clusters that describe the transition to biotrophy. Those clusters enriched in putatively secreted proteins (S), effectors (E), and cell wall modifying enzymes (C) are shown (e.g. SE denotes a cluster is enriched in putatively secreted proteins and effectors). N.B. effectors and cell wall modifying enzymes are also putatively secreted proteins. **B**) For each cluster, the proportion of genes present in each orthologous gene cluster category is shown for the following species *Caenorhabditis elegans, Bursaphelenchus xylophilus, Meloidogyne graminicola, M. hapla, Globodera pallida, G. rostochiensis, Heterodera glycines*, and *H. schachtii*. The dotted line represents the percentage of all DEGs in that orthologous gene cluster category for reference.

The only cluster significantly enriched in CWMEs is J2_10hpi (hypergeometric test, FDR adj. p-value = 5.41e^-10^) consistent with the idea that CWMEs facilitate both entry into, but also movement within, the host (i.e. not specific to J2). While 65% of all putative orthologues of known effectors are contained within five clusters that describe the transition to biotrophy, only one of these (48 hpi) is significantly enriched (hypergeometric test, FDR ad. p-value = 1.08e^-49^) and contains 38% of all putative orthologues of known effectors. The only other cluster enriched for effectors contains exclusively life stages involved in biotrophic interaction with the host (48hpi_12dpifem_12dpimale_24dpi, hypergeometric test, FDR adj. p-value = 1.15e^-10^).

Cross referencing the differential expression clusters that describe the transition to biotrophy with the orthogroup analysis suggests that the transition to biotrophy is generally characterised by a proportional decrease in genes shared among nematodes, and a proportional increase in various orthologous genes cluster categories specific to subsets of plant-parasitic nematodes. On a finer scale, how the distribution of genes across the orthologous gene cluster categories compares between differential gene expression clusters that describe the transition to biotrophy is very informative. To summarise the biggest differences: the beginning of the transition to biotrophy (J2, and J2_10hpi) is characterised by the greatest proportional increases in plant-parasitic nematode-specific genes and cyst and root knot nematode-specific genes, and the greatest proportional decreases in *Heterodera*-specific genes (Figure 4B); the middle of the transition (10 hpi) is characterised by a proportional increase in *Heterodera*-specific genes; and the end of the transition (10hpi_48hpi and to a greater extent 48 hpi) is characterised by a proportional increase in cyst nematode-specific genes and *Heterodera*-specific genes. Considering the functional annotation of genes in these clusters (Figure 4A), we conclude that genes expressed in the J2 are not involved in host-manipulation (unless they are also expressed in another life stage (e.g. J2_10hpi)). Given that the 10 hpi cluster is characterised by a larger proportional increase in *Heterodera-specific* genes than cyst nematode-specific genes, coupled with the fact that the greatest difference between infected and uninfected plant tissue is 10 hpi, these data appear consistent with the idea that a greater degree of specialisation from the parasite is needed to overcome the early host response. Finally, we conclude that *H. schachtii* is a fantastic model for the most agriculturally important cyst nematode, *H. glycines*, because *H. schachtii-specific* genes represent a tiny proportion of those expressed during the transition to biotrophy (generally <5%).

Subsequently, we analysed the five differential expression clusters, of the host and of the parasite, that describe discrete stages of infection (10hpi, 48hpi, 12dpi_male, 12dpi_female, and 24dpi_female; Figure 5). Strikingly, host and parasite genes upregulated at the same time of infection have different, and contrasting, evolutionary histories within their respective phyla. Parasite genes in expression clusters that peak at individual times of infection are generally less well conserved across the phylum Nematoda when compared to all differentially expressed genes in the experiment (Figure 5; Figure S5). In contrast, host genes in expression clusters that peak at those same times of infection are generally more well conserved across the phylum Planta when compared to all differentially expressed genes in the experiment (Figure 5; Figure S5). Although these evolutionary time scales are not directly comparable, these data may suggest that infection-specific gene expression is characterised by “lineage-specific” nematode genes (including, for example, effectors) modulating the expression of “widely conserved” plant genes (including, for example, genes involved in development of the feeding site).

**Figure 5.**
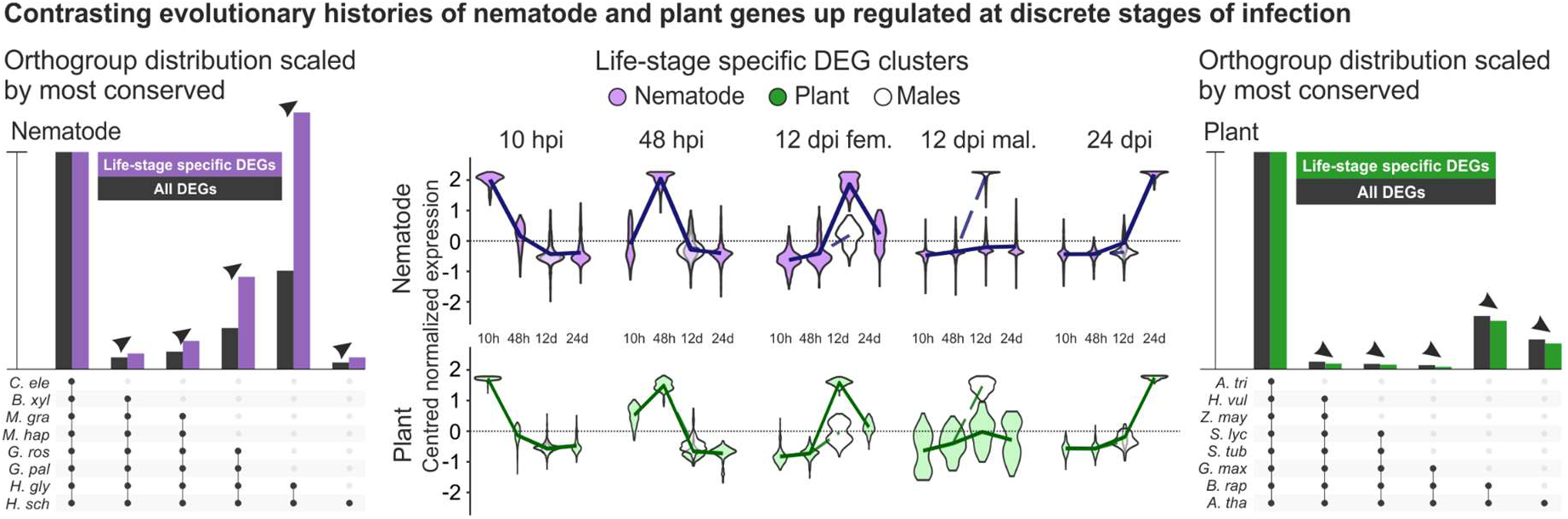
Contrasting evolutionary histories of host and parasite genes deployed at specific times of infection. Differential expression super clusters that describe discrete stages of infection (centre) for either the host (closed green) or the parasite (closed purple) where hpi = hours post infection and dpi = days post infection. Open violins represent males, or plant tissue associated with males. The orthogroup distributions (scaled to the most conserved category) are shown for eight nematode species (left - *Caenorhabditis elegans, Bursaphelenchus xylophilus, Meloidogyne graminicola, M. hapla, Globodera pallida, G. rostochiensis, Heterodera glycines*, and *H. schachtii*) and eight plant species (right - *Amborella trichopoda, Hordeum vulgare, Zea mays, Solanum lycopersicum, S. tuberosum, Glycine max, Brassica rapa* (ssp. *rapa)* and *A. thaliana*). the parasite (left) and the host (right). Arrowheads indicate proportional increase or decrease compared to all differentially expressed genes (DEGs, black).

### Congruent differential expression of metabolic pathways during infection highlights nematode susceptibility genes

To understand the roles of “widely conserved” host genes, and “lineage-specific” parasite genes, that are differentially regulated during biotrophy (Figure 5), we used the KEGG Automatic Annotation Server Ver. 2.1 (https://www.genome.jp/kaas-bin/kaas_main) to annotate metabolic pathways in the host (3,860 KO terms on 9,240 genes) and the parasite (3,479 KO terms on 4,990 genes (Table S8)). Given the partial overlap of KO terms (1,998 in common between host and parasite) we manually examined pathways that were incomplete in the parasite, but complemented by the host.

The vitamin B5 (pantothenate) biosynthetic pathway (M00119) is an exquisite example of congruent differential expression of metabolism involving “widely conserved” host genes and “lineage-specific” parasite genes. The complete pathway is conserved in most plants, and all steps (even when more than one gene encodes a particular function) are upregulated during infection of *A. thaliana* with *H. schachtii* (Figure 6A; Figure S6), with the exception of the last step pantothenate synthetase (AtPANC). The dominant peak of expression in all upregulated cases is at 12 dpi in the syncytia associated with the female. The expression of this pathway in syncytia associated with the 12 dpi male is either indistinguishable from the control, or less highly upregulated than in syncytia associated with the 12 dpi female, and approximately at the same level as in the syncytia associated with the 24 dpi female. While all animals require vitamin B5 in order to complete basal metabolism of proteins, carbohydrates, and lipids, the biosynthesis pathway is considered absent (vitamin B5 is an essential nutrient for animals). Unusually for an animal, the *H. schachtii* genome encodes two genes (*Hsc_gene_23032* and *Hsc_gene_23033*) annotated to carry out only the last step (PANC) of the vitamin B5 biosynthesis pathway (Craig et al. 2009). These putative *H. schachtii PANC* (*HsPANC*) genes are dissimilar in sequence to plant orthologues. Querying the non-redundant nucleotide archive, the most similar sequences to *HsPANC* are from a few animals with obligate interactions with plants (primarily, although not exclusively, plant-parasitic nematodes) and the vast majority from bacteria (Figure 6B). Phylogenetic analyses group the *HsPANC*, along with many other plant-parasitic PANC-like sequences, towards the base of a well-supported clade (100 bootstrap) containing sequences primarily from actinobacteria (the most closely clustered sequence being from *Candidatus Rhodoluna planktonica*, WP_070954084.1, a well characterised but as-yet uncultured organism). Importantly, both *HsPANC* encode several well supported introns and are in the middle of a large 1.3 Mb scaffold surrounded by classical nematode genes, disproving the possibility of contamination (Figure 6C). The most parsimonious explanation for the existence of *PANC* in the *H. schachtii* genome is therefore their acquisition by horizontal gene transfer from bacteria to the last common ancestor of the suborder Hoplolaimina - indicative of an important role in plant-nematode interactions across the group - and subsequent tandem duplication in *H. schachtii* or its progenitor. Remarkably, *HsPANC* complement the upregulation of the first steps in the pathway in the host (and the absence of upregulation of the last step in the pathway in the host): *HsPANCs* are upregulated in the 12 dpi female, to a lesser extent in the 12 dpi male, and to a greater extent in the 24 dpi female (Figure 6A).

**Figure 6.**
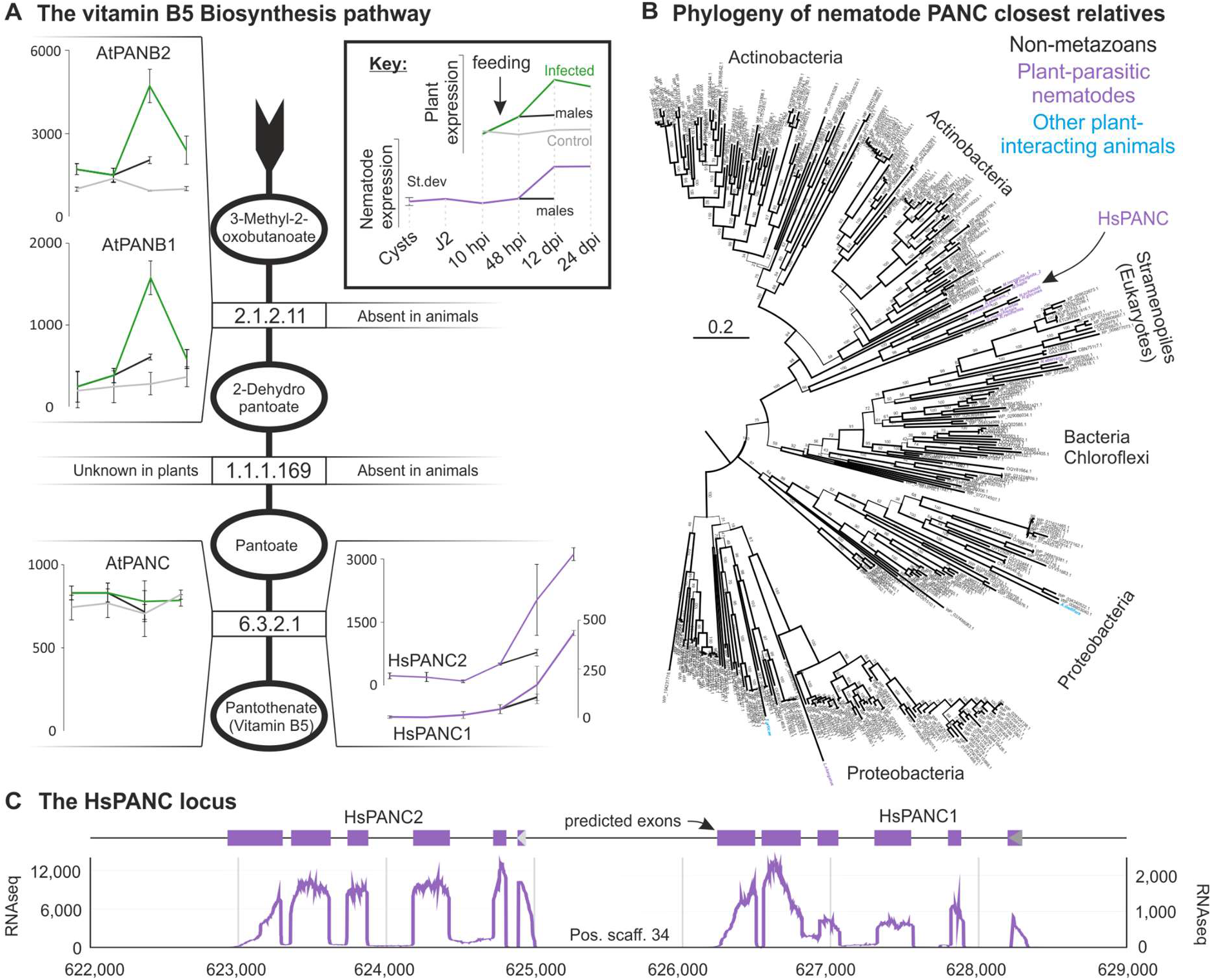
Congruent differential expression of the vitamin B5 biosynthesis pathway between kingdoms is enabled by a horizontal gene transfer from bacteria. **A)** The linear pathway of vitamin B5 biosynthesis. Products/substrates are indicated with circles, enzymatic reactions (and corresponding EC codes) with squares. For each species, the expression profiles of the gene/s annotated with the EC codes at each step are shown (green for host, purple for parasite), if present. **B)** Phylogenetic tree inferred from the protein alignment of the top 50 most similar sequences identified in the NCBI non-redundant database to each of 14 animal putative *PANC* genes. Most *PANC* homologues from plant-parasitic nematodes (purple), including *HsPANC*, group in a single monophyletic sub-clade with sequences from actinobacteria. **C)** The two predicted *H. schachtii PANC* orthologues are adjacent, in the same orientation, and located towards the middle of a 1.3 Mb scaffold. RNAseq read mapping (graph) and intron:exon structure of both genes (5:6, blocks above) are similar, and confirm that they do not arise from contamination.

Taken together, we conclude that the vitamin B5 (pantothenate) biosynthetic pathway is congruently differentially expressed between kingdoms in this interaction. The presence of PANC-like sequences in two other groups of animals outside the plant-parasitic nematodes (*Tetranychus urticae* - a species of plant-feeding mite generally considered to be a pest - XM_015927639.1, and *Apis mellifera* - the western honeybee - XM_001123024.1) is intriguing (given their similarly obligate relationship with plants), but more ambiguous than for the plant-parasitic nematodes. The predicted protein sequences are 52% and 65% identical with the most similar bacterial sequence in NCBI (with much less similar nucleotide sequences), although neither had introns, and the *A. mellifera* gene is on a short scaffold.

These data support the hypothesis that the plant-encoded enzymes of the first part of the vitamin B5 biosynthesis pathway support nematode infection, and may thereby function as nematode susceptibility genes. To investigate whether *AtPANB1* (the penultimate step in the pathway, and the last step upregulated upon infection) plays a role in cyst nematode parasitism, we generated homozygous loss-of-function T-DNA insertion lines for *AtPANB1* (*atpanb1-1*; Figure S7). With the exception of a slight delay in flowering time, *atpanb1-1* was phenotypically indistinguishable from wild-type in the absence of infection: hypocotyl length, root and shoot fresh weight, and siliques sizes, were not altered compared to Col-0 (Figure S8). During infection, however, several marked differences were noted between the mutant and wild-type lines. We found that *atpanb1-1* supported fewer nematodes at 14 dpi (Figure 7A; p-value < 0.0001). Moreover, even those nematodes that were able to infect the mutant are less fit as evidenced by the formation of smaller syncytia (Figure 7B, D, p-value < 0.0001), smaller females (Figure 7C, D, p-value = 0.02), smaller cysts (Figure 7E; p-value = 0.049), and lower average number of eggs per cyst (Figure 7F; p-value = 0.049). Importantly, the reduced susceptibility of *atpanb1-1* to nematodes can be rescued by overexpressing *AtPANB1* under the control of a *35S* promoter in the *atpanb1-1* background (Figure 8).

**Figure 7.**
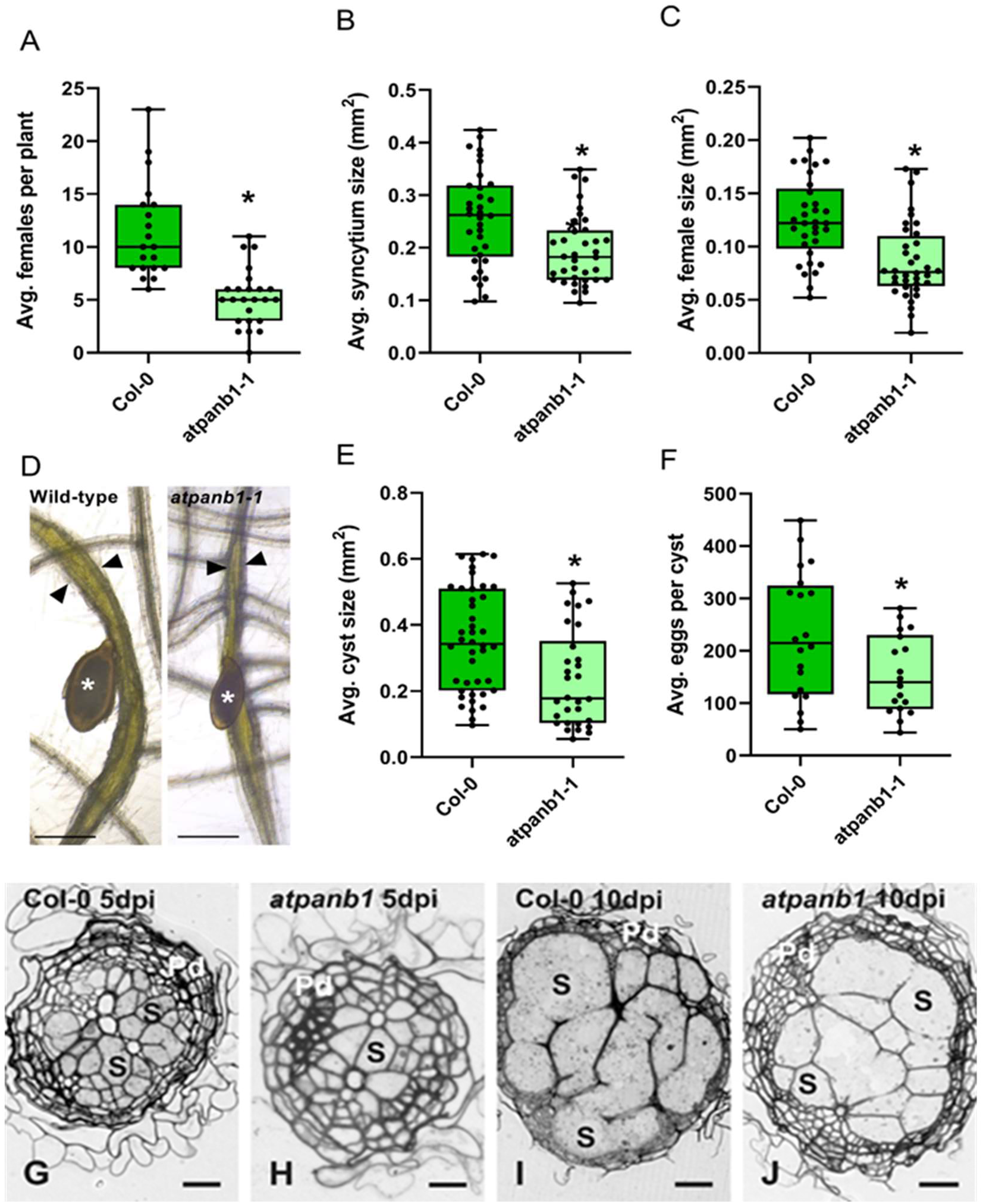
*AtPANB1* is required for cyst nematode parasitism. **A)** Average number of female cyst nematodes present per plant root system at 14 dpi. Approximately 60 - 70 *H. schachtii* infective juveniles were inoculated onto 12-day-old Arabidopsis plants. The number of females per root system was counted at 14 dpi. **B**) Average size of syncytia at 14 dpi. Approximately 25 - 40 syncytia were randomly selected, and their outlines were measured for each independent experiment. **C**) Average size of female nematodes at 14 dpi. Approximately 25 - 40 female nematodes were randomly selected, and their outlines were measured for each independent experiment. **D**) Example images of nematode infection on wild-type and *atpanb1-1* mutants. Arrowheads indicate syncytium boundaries and asterisks indicate female nematodes. **E**) Average size of cysts (20 - 30) repetitions per experiment) at 42 dpi. **F**) Average number of eggs per cyst (15-20) repetitions per experiment) at 42 dpi. Experiments were performed four times independently with the same outcome. Bars represent means ±SE. Data from one experiment is shown. Data were analysed using Student’s t-test. Asterisks indicate significantly different means (95% confidence). **G-J**) Light microscopy images of cross sections taken through the widest part of syncytia induced by *H. schachtii* in Col-0 (**G** and **I**) and *atpanb1-1* (**H** and **J**) roots at 5 (**G** and **H**) and 10 dpi (**I** and **J**). Abbreviations: Pd, periderm; S, syncytium. Scale bars: 20 μm.

**Figure 8.**
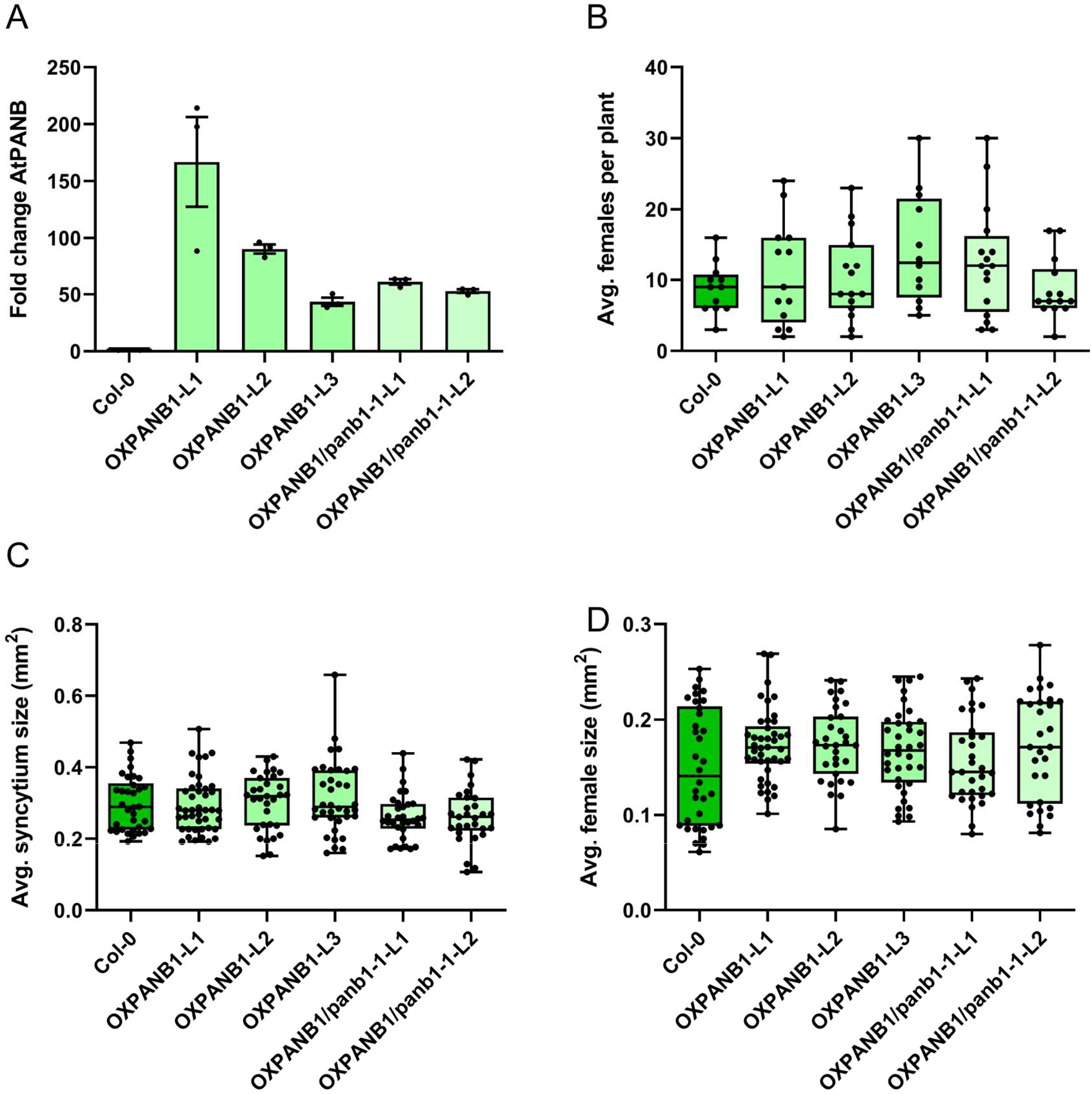
Overexpression of *AtPANB1* restores susceptibility to cyst nematodes. **A)** Quantitative reverse transcription-polymerase chain reaction (qRT-PCR) confirmation of the increase in *AtPANB1* transcript in overexpression lines of the Col-0 *(OXPANB1* lines L1, L2, L3) or *panb1-1* (OXPANB1/panb1-1 lines L1 and L2) background. **B**) Average number of female cyst nematodes present per plant root system at 14 dpi. Approximately 60 - 70 *H. schachtii* infective juveniles were inoculated onto 12-day-old Arabidopsis plants. The number of females per root system was counted at 14 dpi. **C**) Average size of syncytia at 14 dpi. Approximately 25 - 40 syncytia were randomly selected, and their outlines were measured for each independent experiment. **D**) Average size of female nematodes at 14 dpi. Approximately 25 - 40 female nematodes were randomly selected, and their outlines were measured for each independent experiment. Bars represent mean ±SE. **B–D**) Experiments were performed 3 - 6 times independently with the same outcome. Data from one experiment is shown. Data were analysed using a one-way analysis of variance (ANOVA) followed by Tukey’s HSD post-hoc test. No significant difference was detected (95% confidence).

Next, we conducted microscopic examinations of anatomical and ultrastructural features of syncytia induced in roots of *atpanb1-1* plants and compared them with syncytia induced in wild-type Col-0 plants (Figure 7GJ; Figure S9). In both genotypes, syncytia were induced in the vascular cylinder and they were composed of vascular cylinder cells only (Figure 7G-J; Golinowski et al. 1996). In wild-type Col-0, syncytia were surrounded by a continuous layer of periderm (Figure 7G; Figure S9B, D; Holbein et al. 2019), excluding regions next to nematode heads (Figure S9A, C). However, in the case of syncytia induced in *atpanb1-1* roots, the periderm layer was not yet differentiated next to the nematode head (Figure S9E) and syncytia developed poorly at 5 dpi (Figure 7H; Figure S9F). Overall, syncytia induced in *atpanbb1-1* were less well developed than in Col-0 plants as indicated by reduced size and number of cells incorporated into syncytia. This was especially evident at 5 dpi (Figure 7G, H; Figure S9A, B, E, F). At 10 dpi, syncytia induced in *atpanb1* roots were still smaller on cross sections than syncytia induced in wild-type plants, but this difference was less apparent (Figure 7I, J; Figure S9D, H). At the ultrastructural level the organisation of syncytial protoplasts in Col-0 plants was typical for syncytia induced in Arabidopsis roots. Syncytial cytoplasm is marked by a high density of organelles (Figure S9I, J and (Golinowski et al. 1996, Sobczak et al. 1997)). A contrasting situation was found in syncytia induced in *atpanb1-1* roots (Figure S9K, L). At the widest parts of syncytia, some distance from the nematode heads, the syncytial cytoplasm exhibits large regions almost free from plastids, mitochondria and structures of endoplasmic reticulum both at 5 and 10 dpi. These ultrastructural features are unusual, and might indicate that syncytia induced in *atpanb1-1* plants are metabolically less active and therefore less efficient in supporting nematode development leading to/stemming from the formation of smaller and undernourished females.

To investigate whether AtPANB1 plays a similar role in other plant-interacting organisms, we analysed changes in susceptibility of *atpanb1-1* towards the necrotrophic fungus *Botrytis cinerea*, and the beneficial endophyte *Piriformospora indica*. In both cases, no differences were detected between *atpanb1-1* and the wild-type control (Figure S10A, B). Next, we investigated whether the decrease in susceptibility of *atpanb1-1* to cyst nematodes is due to changes in classical immune responses. To this end, we tested ROS production, a hallmark of basal defence in plants, upon treatment of an immunogenic peptide, flg22. We did not observe any changes in ROS burst, both over time or cumulative (Figure S10C). Taken together, these data suggest that *AtPANB1*, the last step in the pathway upregulated during infection, is a specific non-immunity related nematode susceptibility gene. Given that this pathway is linear, and that all previous steps are similarly upregulated, the implication is a series of nematode susceptibility genes for future study. The wide conservation of *PANC* in plant-parasitic nematodes (Figure 6B) extends this intriguing possibility to most other plant-parasitic nematodes of global agricultural importance (J. T. Jones et al. 2013).

### Trans-kingdom compartmentalization of the vitamin B5 pathway

In contrast to *AtPANB*, *AtPANC* is neither upregulated in syncytium nor plays a role in cyst nematode infection: loss-of-function homozygous mutants for *AtPANC1* (Figure 6; Figure S6, S7) are indistinguishable from wild-type plants during infection (Figure 9A-C). Complementing this lack of upregulation, the expression of both *HsPANC1 and HsPANC2* is not only increased after onset of parasitism (48 hpi) but remains high as the nematode proceeds through the rest of its life cycle, pointing to a functional role for *HsPANC* in parasitism (Figure 6A). To test this hypothesis experimentally, we used *in vitro* RNAi (Figure 9D), which caused a significant decrease in expression of *HsPANC* and disrupted the nematode’s ability to infect wild-type Col-0 (Figure 9E). The average sizes of the females and syncytia were also significantly reduced on Col-0 infected with nematodes showing a reduced expression of *HsPANC* (Figure 9F,G). Importantly, HsPANC functions in the nematode. *HsPANC* does not encode an N-terminal secretion signal, and *in situ* hybridization of digoxigenin labelled probes to *HsPANC* mRNA reveals expression throughout the nematode body with a particularly strong signal in the intestinal region of the nematode (Figure 9H). Taken together, we conclude that the contribution of HsPANC to parasitism success is due to its role in the nematode.

**Figure 9.**
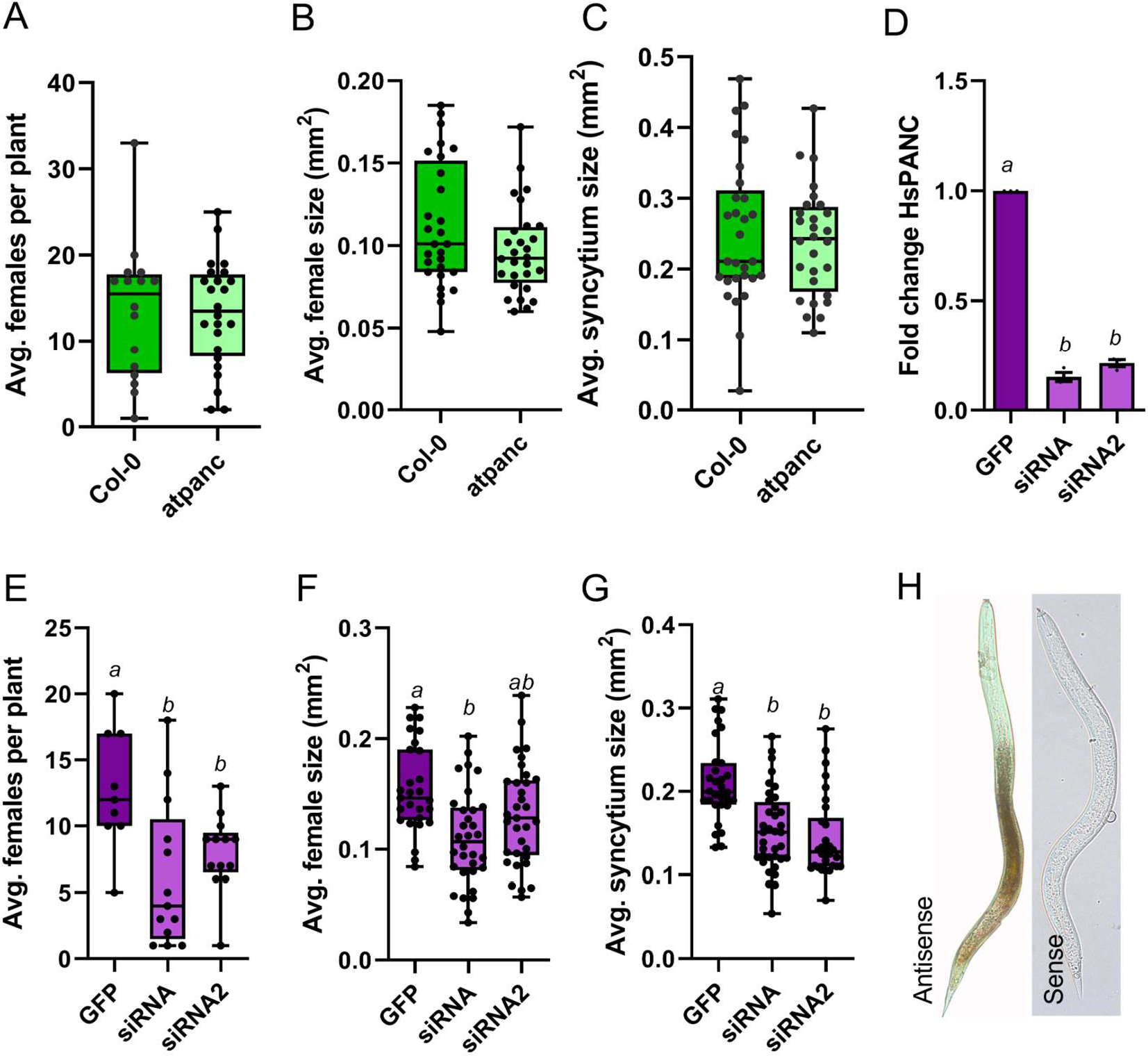
HsPANC, but not AtPANC, is required for cyst nematode parasitism. **A**) Average number of female cyst nematodes present per plant root system at 14 dpi. Approximately 60 - 70 *H. schachtii* infective juveniles were inoculated onto 12-day-old Arabidopsis plants. The number of females per root system was counted at 14 dpi. **B**) Average size of syncytia at 14 dpi. Approximately 25 - 40 syncytia were randomly selected, and their outlines were measured for each independent experiment. **C**) Average size of females at 14 dpi. Approximately 25 - 40 female nematodes were randomly selected, and their outlines were measured for each independent experiment. **D**) Change in transcript abundance of *HsPANC* gene in J2 nematodes soaked in siRNA targeting *HsPANC* or *GFP*. **E)** Average number of females present per Col-0 root system infected with J2 soaked in siRNA targeting *HsPANC* or *GFP* at 14 dpi. Approximately 60 - 70 *H. schachtii* infective juveniles were inoculated onto 12-day-old Arabidopsis plants. The number of females per root system was counted at 14 dpi. **F)** Average size of females in Col-0 infected with J2 soaked in siRNA targeting *HsPANC* or *GFP* at 14 dpi. Approximately 25–40 female nematodes were randomly selected, and their outlines were measured for each independent experiment. **G**) Average size of syncytia in Col-0 infected with J2 soaked in siRNA targeting *HsPANC* or *GFP* at 14 dpi. Approximately 25 - 40 syncytia were randomly selected, and their outlines were measured for each independent experiment. **H**) *In situ* hybridization of the *HsPANC* gene in J2 of cyst nematode. **A-C)** Experiments were performed three times independently with the same outcome. Data from one experiment is shown. Data were analysed using Student’s t-test. No significant difference was detected (95% confidence). **D–G**) Experiments were performed three times independently with the same outcome. Data from one experiment is shown. Bars represent mean ± SE. Data were analysed using a one-way analysis of variance (ANOVA) followed by Tukey’s HSD post-hoc test. Different letters indicate significantly different means (95% confidence).

To understand how HsPANC contributes to parasitism success in the nematode, we tested whether *HsPANCs* encode a functional enzyme. *HsPANC1* cDNA was cloned into PUC18, and was tested for its ability to complement *panb* mutant from *E. coli* (AT1371). The *E. coli PANC* gene served as a positive control. All strains grew well in the presence of pantothenate in the growth medium. In absence of exogenous pantothenate, the *panc* mutant containing the empty vector did not grow. However, the strains transformed with either bacterial or nematode *PANC* grew well with no significant differences detected between strains (Figure 10). We therefore concluded that *HsPANC* encodes a bona fide PANC.

**Figure 10.**
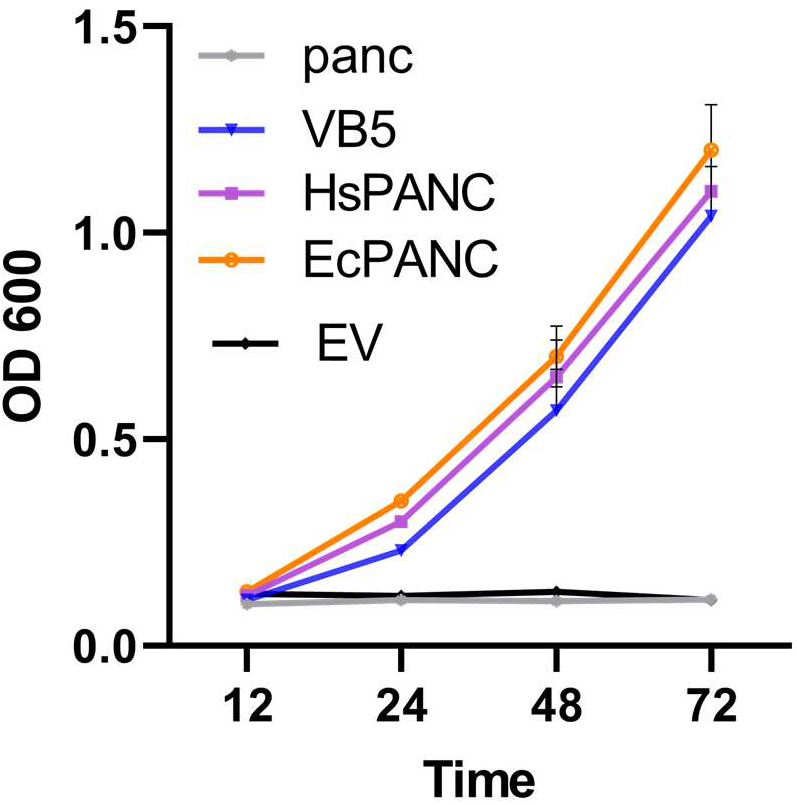
*HsPANC* encodes a functional PANC. Functional complementation of PANC-deficient *E. coli* strain (AT1371) (panc) and containing the empty plasmid (EV), exogenous pantothenate (VB5), nematode PANC (HsPANC), and the bacterial PANC (EcPANC). Bars represent mean ±SE from three independent experiments.

The presence of *PANC* homologs in the cyst nematode’s genome indicates an evolutionary benefit, but the molecular fundamentals of this peculiar separation remain elusive. Previous studies show that plant PANC is subjected to uncompetitive substrate inhibition at higher pantoate concentrations (Genschel et al. 1999). Similarly, the activity of PANC is also inhibited by products, and in particular by accumulation of pantothenate (Ronconi 2006). Based on these observations, we propose a hypothesis (Figure 11) that compartmentalization of vitamin B5 pathway between plants and nematodes helps to avoid feedback/feed-forward inhibition and ensures a consistent supply of vitamin B5 to rapidly developing nematodes. Clarifying further details of the nematode vitamin metabolism, including details of the activity of HsPANC and AtPANC may aid in the development of nematode-resistant germplasm through the targeted deletion of “susceptibility” genes.

**Figure 11.**
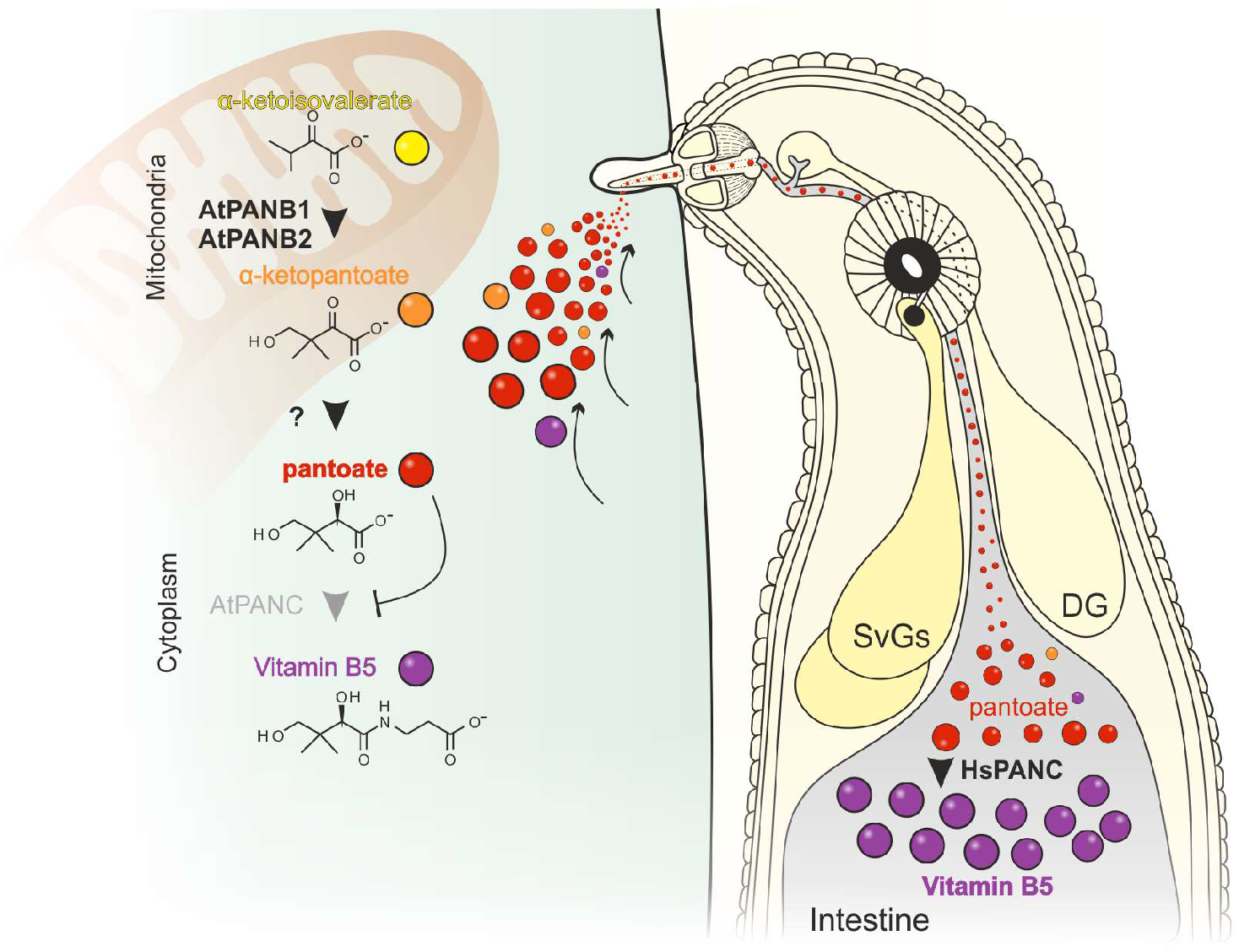
A model for the trans-kingdom synthesis of vitamin B5. Left, plant cell. Right, parasitic nematode. The last two steps of the vitamin B5 biosynthesis pathway are indicated (bold names indicate upregulated during parasitism), including their sub-cellular localisation. Products/substrates are represented as globes.

## Conclusions

- *H. schachtii* has the largest genome, and second largest exome, of any cyst nematode analysed to date.
- Substantial segmental duplication contributed to the large gene number, started at some point in the Heterodera lineage after the split from Globodera, was still active in the *H. schachtii* lineage after the split from *H. glycines*, and has acted on old and new genetic capital.
- A reference trans-kingdom, life stage-specific, transcriptome of *H. schachtii* and *A. thaliana* reveals that host and parasite genes upregulated at the same time of infection have different, and contrasting, evolutionary histories within their respective phyla.
- Congruent differential expression analysis of metabolic pathways highlights nematode susceptibility genes.
- The vitamin B5 biosynthesis pathway of the plant-nematode hologenome starts in the plant, ends in the nematode, and is putatively enabled by a horizontal gene transfer from a bacterium.
- Transkingdom compartmentalization of the vitamin B5 biosynthetic pathway, ensures a consistent supply of vitamin B5 to the parasite by avoiding feedback/-forward inhibition.

## Methods

### Flow cytometry

Approximately 50 μl of compacted live *H. schachtii* J2s (Bonn population) in a 1.5 ml eppendorf tube were resuspended in 10 μl of extraction buffer (PBS with 0.01% Triton X) and disrupted with a micro pestle. Lysed nematodes were resuspended in 500 μl of extraction buffer and the debris filtered through 100 μm and then 30 μm filters (CellTrics). Up to 250 μl of the filtrate was added to 1 ml of staining buffer (100 mM Tris (pH 7.4), 150 mM NaCl, 1 mM CaCl_2_, 0.5 mM MgCl_2_, 2 μg ml^-1^ propidium iodide, and 0.18 mg ml^-1^ RNAse A) and incubated on ice in the dark for 1 h. Stained nuclei were counted on an Accuri C6 flow cytometer (BD Biosciences-US). For *A. thaliana*, 1-2 leaves were removed and placed on a Petri dish, immersed in 250-500 μl of extraction buffer. The leaf tissue was finely cut with a razor blade. The remaining liquid and leaf debris was filtered, stained, and counted as for nematode samples. Flow cytometry data were analysed with FlowJo.

### Trans-kingdom transcriptome sampling, sequencing, and analysis

Seeds of *Arabidopsis thaliana* ecotype Columbia-0 (Col-0) were surface sterilized by washing in 70% (v/v) ethanol for 5 min followed by washing with 2% (w/v) sodium hypochlorite for 3 min. Seeds were rinsed twice with sterile water and were dried on sterile Whatman filter paper before planting. Plants were grown in Petri dishes in agar supplemented with modified Knop’s nutrient medium at 23 °C under long day conditions with 16 h light and 8 h darkness. Second stage juveniles were harvested from nematode stock culture, which has been maintained on mustard roots under sterile conditions for 20 years. Twelve-days-old plants were inoculated with 60 - 70 surface-sterilized juveniles. Small root segments containing nematodes were marked under a stereo microscope. The infected area including nematodes was then hand-dissected and transferred to liquid nitrogen. Several hundred segments were collected for each biological replicate. Uninfected root segments adjacent to the infected area were used as a control.

Frozen tissue was lysed to a fine powder with a tissue lyser at 30 Hz (2 x 2 min). RNA was extracted from frozen powder using the RNeasy Plant Mini Kit (Qiagen) following the manufacturer’s instructions and using both the optional shredder columns and on column DNA digestion. RNA quality was assessed using Bioanalyzer. Library construction (150 bp paired end stranded RNAseq) and sequencing was carried out by Novogene (Bioproject: PRJNA722882). Additional sequencing of the early mixed time points (10 h and 48 h post infection) was carried out to provide enough coverage of the nematode exome (Figure 1).

### Genome extraction, sequencing, and assembly

High molecular weight DNA was extracted from pools of approximately one million frozen *H. schachtii* J2s (Bonn population) using the phenol/chloroform/isoamyl alcohol extraction method. In brief: pelleted juveniles (in a 1.5 ml eppendorf) were resuspended in 160 μl of extraction buffer (0.1 M Tris (pH 8.0), 0.5 M NaCl, 50 mM EDTA, 1% (w/v) SDS). The pellet was ground using a micro pestle for less than 30 s and until the pellet thawed, and 40 μl of proteinase K (20 mg ml^-1^) was added. The juveniles were incubated at 55 °C for 24 h, mixing gently but often. RNAse A (10 μl of 10 mg ml^-1^) was added, incubated at room temperature for 10 min, and mixed with equal volumes of phenol/chloroform/isoamyl alcohol solution (25:24:1). Phenol/chloroform extraction, chloroform back extraction, and ethanol precipitation were performed according to the PacBio protocol (https://www.pacb.com/wp-content/uploads/2015/09/SharedProtocol-Extracting-DNA-usinig-Phenol-Chloroform.pdf).

Two PacBio libraries were prepared from extracted DNA. First, a 25 kb library (DNA was fragmented to 25 kb, size selection of the final library was performed using BluePippin at 10 kb cut-off), and subsequently a 35 kb library (DNA was fragmented to 35 kb, size selection was performed with BluePippin at 15 kb cut-off). Each library was sequenced on Pacific Biosciences Sequel instrument using Sequel Polymerase v2.1, SMRT cells v2 LR and Sequencing chemistry v2.1. Loading was performed by diffusion, in total 4 SMRT cells were used (movie time: 1200 min). Sequencing was provided by the Norwegian Sequencing Centre (www.sequencing.uio.no), a national technology platform hosted by the University of Oslo and supported by the “Functional Genomics” and “Infrastructure” programs of the Research Council of Norway and the Southeastern Regional Health Authorities. In addition, one 150 bp paired end (350 bp insert size) Illumina library was prepared from the same DNA (NEBNext Ultra II DNA Library Prep Kit for Illumina) and sequenced to approximately 140 fold coverage (24.6 Gb) using the service provided by NovoGene (GenBank Accession PRJNA722882).

All PacBio subreads (Bioproject: PRJNA722882) were error corrected using Canu snapshot v1.7 +137 changes (r8829 73d5caa1b1087b65f7853ecbebc1bb1dcbd1bc14) (Koran et al. 2017), with the following additional parameters: corOutCoverage=300 corMhapSensitivity=normal. This allowed all the PacBio data to be used in the error correction phase instead of 40X, resulting in 1,013,782 corrected reads. Corrected reads were assembled using wtdbg2 Version: 2.1 - 20181007 (Ruan and Li 2020). A variety of parameters were tested, and each assembly was assessed for assembly size, BUSCO (Simão et al. 2015) version 1.1b using Eukaryota data set for completeness score (pre-trained on *G. pallida* Augustus version 3.2.1 models (Stanke et al. 2006)), and RNAseq mapping STAR (Dobin et al. 2013). The final assembly used the following parameters: -L 5000 -p 19 -AS2. CEGMA (Parra, Bradnam, and Korf 2007) was also used to quantify the completeness of assemblies based on a set of core eukaryotic genes.

The assembly was subjected to BlobTools version 1.0 analysis (Laetsch and Blaxter 2017) to identify and remove putative contaminant contigs. Briefly, reads were mapped to the assembly using minialign version 0.5.2 (https://github.com/ocxtal/minialign) to determine the coverage of the assembled contigs. The contigs were compared to the GenBank nt database using BLASTn (reporting taxonomic information). Contigs were then classified based on the weight of the BLAST hits. Contigs identified as fungal, bacterial, plant or viral were removed. Thus, yielding a contamination free unpolished contig level assembly.

Haplotigs and either extremely high or extremely low coverage contigs were removed (using purge haplotigs (a-70)) (Roach, Schmidt, and Borneman 2018). Remaining contigs were upgraded using FinisherSC (Lam et al. 2015) and scaffolded using iterative rounds of SSPACE_longreads (Boetzer and Pirovano 2014) and gapFinisher (Kammonen et al. 2019): Round 1 (error corrected reads, -k 1 -o 1000 -l 10 -g 500); round 2 (error corrected reads,-k 1 -o 1000 -l 5); and finally round 3 (raw reads, -k 1 -o 1000 -l 30). Scaffolds were then polished with three rounds of arrow (raw PacBio reads, smrttools-release_6.0.0.47835, in --diploid mode), and five rounds of Pilon (raw PacBio reads mapped with minimap2, 150 bp Illumina read pairs mapped with BWA-mem). This gave rise to assembly version 1.1.

During initial phases of gene prediction on assembly version 1.1, it was noted when mapping RNAseq to the assembly that some SNPs in genes (often, but not always, A|G polymorphisms) were miscalled as adjacent indels (therefore altering the translation frame and negatively affecting gene calls). Mapped RNAseq was therefore used to create a variant call file (Pilon), which was parsed with a custom python script to identify all indels that were up to 4 bp apart (1,451). All instances were inspected in a genome browser and corrected, giving rise to the final manually finished assembly version 1.2 (GenBank Accession JAHGVF000000000). Scripts, commands, and config files are available at: https://github.com/peterthorpe5/Heterodera_schachtii_genome_assembly and https://github.com/sebastianevda/H.schachtii_genome/assembly_and_manual_finishing_scripts).

### Gene prediction and annotation

Hard and soft genome masking was performed as described in (Eves-van den Akker et al. 2016; Thorpe et al. 2018). Briefly, Repeatmodeler (version DEV) was used to identify repetitive regions. The resulting identified repetitive elements were masked using RepeatMasker along with RepBaseRepeatMaskerEdition-20170127 models. To additionally identify transposons, TransposonPSI version 08222010 (Haas 2007) and LTRharvest version 1.5.9 (Ellinghaus, Kurtz, and Willhoeft 2008) from Genometools (Gremme, Steinbiss, and Kurtz 2013) was used. All RNAseq reads were remapped to the soft masked genome, separated by strand, and used to provide stranded hints for Braker2 gene prediction (-f 2000, allowing Augustus to train itself for both exon and utr models). After considerable optimisation of extrinsic config files for exon and UTR prediction (available at https://github.com/sebastianevda/H.schachtii_genome/Gene_predictions), an extremely high quality set of annotations resulted (termed annotation version 1). The version 1 predicted proteins were annotated with DIAMOND-BLASTp (version v0.9.24.125) (Buchfink, Xie, and Huson 2015) searched against GenBank NR database. The resulting .xml file was imported into BLAST2GO version 5 (Conesa et al. 2005). BLAST2GO was used to annotate the gene models using GO version May 9^th^ 2019 and Interproscan (Quevillon et al. 2005). Gene duplication analyses were performed using the similarity searches from DIAMOND-BlastP (1e-10) with MCSanX toolkit (Wang et al. 2012). PFAM domains in protein coding sequences were predicted using hmmsearch version 3.2.1 (Eddy 2011) and the Pfam-A.hmm definitions release 30 (http://ftp.ebi.ac.uk/pub/databases/Pfam/). Signal peptide and transmembrane prediction was performed using signalP 4.0 (Petersen et al. 2011) and Phobius version 1.01 (Käll, Krogh, and Sonnhammer 2004).

### Effector identification

Genes related to previously reported effectors in *G. rostochiensis* supported by gland cell expression were identified in the *H. schachtii* genome as previously (Eves-van den Akker et al. 2016). In brief, an inclusive list of effectors was generated by sequence similarity alone (either BLAST (e-value < 1e-5) or using HMM searches (for glutathione synthetase-like effectors (Lilley et al. 2018) (PF03199.10 and PF03917.12) and SPRY-domain (PF00622.23) containing effectors (Rehman et al. 2009) (HMMsearch with gathering threshold cut off)). Putative effector families were aligned and manually curated based on expert knowledge of effector characteristics (e.g. low scoring sequences without the appropriate effector domains for a particular effector family were removed). Gene models were manually inspected and re-annotated using Apollo where the gene prediction had failed to correctly capture the intron:exon structure of known effectors. Finally, putative effectors were cross referenced with putative secreted proteins (i.e. encode a signal peptide but do not encode a transmembrane domain) to produce a high-confidence list of effectors (Table S6).

### Horizontal gene transfer and phylogenetics

We used BLAST+ to perform a similarity search of *H. schachtii* proteins against the nr database. Alien index (AI) was calculated with the Alienness web server (Rancurel, Legrand, and Danchin 2017). We then used the AvP software (https://github.com/GDKO/AvP) to perform the HGT analyses. Proteins with AI above 10 were selected and grouped based on the percentage of shared BLAST hits (70%) using single linkage clustering. For each group we extracted the FASTA sequences of the BLAST hits from the NR database. Sequences for each group were aligned using MAFFT (Katoh and Standley 2013) and FastTree (Price, Dehal, and Arkin 2010) was used for phylogenetic inference. In total 540 proteins (containing alternative transcripts) were analysed, with 349 being confirmed as HGT candidates (222 of them had only hits from Non Metazoan taxa), 97 proteins had a non-conclusive topology, and in 94 proteins the hypothesis of being an HGT candidate was rejected. For the 127 proteins confirmed as HGT candidates that contained both Metazoan and non-Metazoan proteins, we performed a constrained phylogenetic inference enforcing the *H. schachtii* protein to form a monophyletic group with the Metazoan proteins. We then performed tree topology tests with IQ-TREE 2 (Minh et al. 2020) to check whether the constrained phylogenetic tree was significantly worse than the unconstrained phylogenetic tree. This analysis showed that in 30 cases both trees were equally likely, bringing down the total number of potential HGT candidates to 319 derived from 263 genes.

Nematode and other animal *PanC* sequences were collected from NCBI and wormbase parasite (Howe et al. 2017) using the *H. schachtii PanC* as a query in BLAST. Probable contaminants were removed based on nucleotide identity >75% to non-Metazoans on NCBI and the absence of introns (the majority of non-nematode sequences were removed). Each remaining non-Metazoan *PanC* was used to identify the top 50 most similar sequences in NCBI using BLAST. A non-redundant list of subjects was aligned to all queries using MUSCLE, and refined (Edgar 2004). The alignment was trimmed using Trimal (-gappyout, (Capella-Gutiérrez, Silla-Martínez, and Gabaldón 2009)). Model selection (LG+F+I+G4) and phylogenetic inference (1000 bootstraps) was carried out using IQ-TREE 2 (Minh et al. 2020)). The phylogenetic tree was midpoint rerooted, and formatted using Figtree1.4.

### RNAseq read mapping, normalisation, differential gene expression analysis and clustering

RNAseq reads are available under Bioproject: PRJNA722882. All reads were analysed with FastQC v0.11.8 (Andrews and Others 2010) and trimmed using BBduk v38.34 (https://sourceforge.net/projects/bbmap/). Only the reads with a minimum Phred Quality Score (Ewing et al. 1998) of 30 were kept. Based on the FastQC analysis, 10 nucleotides at the 5’ end of the reads were removed. The reads were also trimmed for the presence of adapters and only reads with a minimum length of 75 bp were used. The trimmed reads from each library were mapped onto the *A. thaliana* TAIR10 genome assembly combined with the *H. schachtii* 1.2 genome assembly using STAR (Dobin et al. 2013). The counting was performed only on the uniquely mapping reads using htseq-count part of HTseq v0.12.4 (Anders, Pyl, and Huber 2014). The count tables were loaded in R v3.5.2 (Computing and Others 2013) and normalized with DESeq2 v1.22.2 (Love, Huber, and Anders 2014). The clustering of each biological replicate was visualized by a Principal Component Analysis using the plotPCA function. Differentially expressed genes were identified using the DESeq2 v1.22.2 package following pairwise comparison between all samples (|log2 fold change| > 0.5 and adjusted p-value ≤0.01). The normalized expression of the differentially expressed genes were loaded then into MATLAB v9.6 R2019a and clustered using the tcap2 plugin (Kiddle et al. 2010). The clustering tables were then loaded and analysed in R using the tidyverse package v1.2.1 (Wickham et al. 2019). Differential expression clusters with different magnitudes but similar profiles were manually grouped into 30 and 29 biologically relevant expression superclusters for the host and parasite respectively. Visualizations were obtained using the R package ggplot2 v3.1.0. All normalized expression data are available in Tables S3 and S4.

### OrthoMCl analyses

Orthogroup analyses were precomputed and already available for plants and nematodes. For plants, we focused on a subset of eight plant species (*Amborella trichopoda* (outgroup), the monocots *Hordeum vulgare* and *Zea mays*, the solanaceous *Solanum lycopersicum* and *S. tuberosum*, the fabaceous *Glycine max*, and the brassicaceous *Brassica* rapa (ssp. *rapa)* and *A. thaliana* - GreenPhyl v5 - MCL matrix for level 4 clusters (Guignon et al. 2021)). Counts per species per orthogroup were generated using a custom python script (https://github.com/sebastianevda/H.schachtii_genome/tree/main/MCL_parsing). Orthogroups, or subsets of orthogroups that contain genes of interest (e.g. genes in a particular expression cluster), were visualised using UpSetR R package. For nematodes, we focused on a subset of eight nematodes species (the free living nematode *Caenorhabditis elegans* (outgroup), the pine wilt nematode *Bursaphelenchus xylophilus*, the root-knot nematodes *Meloidogyne graminicola* and *M. hapla*, the potato cyst nematodes *Globodera pallida* and *G. rostochiensis*, and the *Heterodera* species, *H. glycines* and *H. schachtii* (Grynberg et al. 2020)).

### KEGG mapping

The KEGG Automatic Annotation Server Ver. 2.1 (https://www.genome.jp/kaas-bin/kaas_main) was used to annotate metabolic processes in the host and the parasite using the bi-directional best hit strategy (using the default dataset and parameters with the following exception: minimum bit score increased to 100, Table S8). The KEGG reconstruct server was used to populate existing pathways with metabolic process information.

### Nematode infection assays

Arabidopsis plants were grown in Petri dishes containing agar supplemented with modified Knop’s nutrient medium under conditions described above. The nematode infection assays were performed as described previously (Radakovic et al. 2018). Briefly, 60 - 70 J2s were inoculated to the surface of agar medium containing 12-days-old Arabidopsis plants. The numbers of female nematodes per plant were counted at 14 dpi. Each experiment included 15 - 20 plants per genotype. The syncytia or female nematodes were outlined at 14 dpi using a Leica dissecting (DM2000, Leica Microsystems, Germany), and the area was calculated using LAS software (Leica Microsystems). Approximately 25 - 40 females and associated syncytia were outlined per genotype for each experiment. The cysts were outlined at 42 dpi using LAS software (Leica Microsystems) and the area was calculated as described above. Approximately 20 - 30 cysts were outlined per genotype for each experiment. The number of eggs per cyst was counted by crushing cysts from single plants into the 1 ml of 6% (w/v) NaOCl at 42 dpi. Each assay included 15 - 20 plants per genotype. All infection assays were repeated a minimum of three times.

### *In situ* hybridization

Using *H. schachtii* cDNA as template, digoxigenin (DIG)-labelled probes complementary to a target sequence in *HsPANC1* were amplified in asymmetric PCR. The PCR was performed with single sense (negative control) or antisense primers in presence of DIG-labelled deoxynucleoside triphosphates (dNTPs) (Roche). Freshly hatched J2s were fixed and hybridized with the probes following the protocol of de Boer et al. (1998). The hybridized nematodes then were examined using Leica DMI2000 compound microscope. Primers are listed in Table S9.

### Light and electron microscopy

Plants of Col-0 and *atpanb1-1* were grown in Petri dishes and inoculated with J2s of beet cyst nematode as described above. Samples consisting of root pieces containing syncytia and attached juveniles were collected 5 and 10 dpi. They were processed for light and transmission electron microscopy as described previously (Golinowski et al. 1996; Sobczak et al. 1997; Siddique et al. 2014).

### Generation of overexpression and complementation lines

Full-length coding sequence of *AtPANB1* (AT2G46110) was amplified from cDNA synthesized from RNA isolated from 12-days-old Arabidopsis plants. The amplified sequence was cloned into Gateway cloning vector pDONR207 (Invitrogen, USA) using the Gateway^®^ BP Clonase II Enzyme mix (Invitrogen, USA). The cloned fragments were verified through sequencing and transferred via Gateway recombination into the pMDC32 vector under the control of 35S promoter. The verified constructs were introduced into *Agrobacterium tumefaciens* strain GV3101, which was used for the transformation of 4-to 6-week-old Col-0 or *atpanb1-1* mutant plants by the floral dip method (Clough and Bent 1998). After drying of plants, seeds (T0) were harvested and sterilized before being sown on Knop medium supplemented with 25 ug ml^-1^ hygromycin. Transformants were selected to produce homozygous plants. 2 - 3 independent homozygous lines with the highest upregulation were selected for further studies.

### Gene silencing

We identified two 21-nt long target sequences in *HsPANC1* mRNA that begin with an AA dinucleotide. Sense and antisense oligonucleotides for the 21-nt target sequences were obtained in which the Us were replaced with Ts. The 8-nt long sequence (5’-CCTGTCTC-3’) complementary to the T7 promoter primer were added to the 3’ ends of both the sense and antisense oligonucleotides. Afterwards, dsRNA was synthesized from the sense and antisense oligonucleotides using Silencer^®^ siRNA Construction Kit (Cat. Nr. AM1620) according to the manufacturer’s instructions. The siRNA targeting *eGFP* was used as a control. Approximately 2,000 nematodes were incubated in a dsRNA solution containing siRNA, 3 mM spermidine, and 50 mM octopamine. After 24 h incubation, nematodes were surface-sterilized in 0.05% (w/v) HgCl_2_, and infected on 12-day-old Col-0 Arabidopsis plants. Each plant was inoculated with 60 - 70 J2s. Approximately, 300 - 500 J2s were used for RT-qPCR as described below.

### Real-time quantitative PCR

RNA was extracted using a Nucleospin RNA XS (Macherey-Nagel, Germany) kit according to the manufacturer’s protocol. RNA was treated with DNase1 using a DNA-free™ DNA Removal Kit (Ambion) to remove contaminating DNA. cDNA was synthesized using a High Capacity cDNA Reverse Transcription Kit (Life-Technologies, cat. no. 4368814) according to the manufacturer’s instructions. Transcript abundance was measured using the StepOnePlus Real-Time PCR System (Applied Biosynthesis, USA). Each sample contained 10 μl of Fast SYBR Green qPCR Master Mix with uracil-DNA, glycosylase, and 6-carboxy-x-rhodamine (Invitrogen), 2 mM MgCl_2_, 0.5 μl each of forward and reverse primers (10 μM), 2 μl of complementary DNA (cDNA), and water in 20 μl total reaction volume. Data shown are an average of three independent biological replicates. Each biological replicate consisted of 2 - 3 technical replicates. *18S* was used as an internal control except for nematode samples. For nematode samples, actin was used as an internal control. cDNA was diluted 1:100 for *18S* amplification. Relative expression was calculated by Pfaffl’s method (2001) where the expression of the target gene was normalized to an internal control to calculate the fold change. All primers are listed in Table S9.

### Genotyping and expression analysis of knock-out mutants

T-DNA insertion mutants were ordered from the Nottingham Stock Centre (*atpanb1-1*, SALK_18243C; *atpanc*, Salk_101909). The homozygous lines were checked for lack of expression through RT-PCR using primers listed in Table S9.

### Oxidative burst assay

The measurement of ROS production was carried out via a luminol-based method as described previously (Mendy et al. 2017). Briefly, leaf disks measuring 0.5 cm in diameter were cut from 12-days-old Arabidopsis plants and incubated with water for 12 h. After incubation, each leaf disk was transferred to a well in a 96-well plate containing 15 μl of 20 μg ml^-1^ horseradish peroxidase and 35 μl of 10 mM 8-amino-5 chloro-2,3-dihydro-7phenyl(3,4-d) pyridazine sodium salt (L-012, Wako Chemicals). Next, 50 μl of either flg22 or water was added to individual wells. Light emission was measured as relative light units in a 96-well luminometer (Mithras LB 940; Berthold Technologies) over 60 min and analysed using instrument software and Microsoft Office Excel. The experiments were performed in eight biological replicates.

### Pathogenicity assays

The plant susceptibility to *Botrytis cinerea* was evaluated via a modified protocol according to Lozano-Torres et al. (2014). Briefly, 5 μl drops of conidial suspensions (5 x 10^5^) were placed on the leaves of 4-week-old Arabidopsis plants, grown on the soil in the greenhouse. After inoculation, plants were placed in the dark at 20 °C and 100% relative humidity for 3 days. Next, leaves containing necrotic lesions were cut and the necrotic area was measured as described previously. *Piriformospora indica* assays were performed according to Daneshkhah et al. (2013). Briefly, 5 mm *P. indica* mycelium plugs, grown at 28 °C on CM medium, were inoculated to 7-days-old Arabidopsis seedlings grown on MS medium (Duchefa Biochemie, The Netherlands). Fresh roots and shoots weights were measured 7 days after inoculation.

### Bacterial complementation assay

Functional complementation analysis was performed in *E. coli*ΔpanC mutants AT1371 (auxotroph for vitamin B5). Full length ORFs of *AtPANC, EcPANC*, and *HsPANC* were synthesized by Genewiz LLC and cloned into pUC18 vectors. The amplified fragments were cloned in frame with the *lacZ α* gene (encoding α peptide of β-galactosidase) under the control of an inducible *lacZ* promoter. All the constructs were transferred into the competent cells of AT1371 by heat shock method. Transformants carrying pUC18 empty vector were used as a negative control. The transformants were grown overnight on LB agar medium containing Ampicillin (100 μg ml^-1^). Positive clones were selected by colony PCR. Growth curve assay was performed by growing transformants overnight in LB medium. A loop of cells was added to M63 minimal media broth devoid of D-pantothenic acid but supplemented with Ampicillin (100 μg ml^-1^) and IPTG (1 mM). Cultures were grown at 37 °C at 120 rpm for 24 - 72 h. Growth of the bacteria was assayed in terms of optical density (OD600nm) of the cultures.

## Data availability

This Whole Genome Shotgun project has been deposited at DDBJ/ENA/GenBank under the accession JAHGVF000000000. The version described in this paper is version JAHGVF010000000. Code and scripts used in the manuscript are available here: https://github.com/sebastianevda/H.schachtii_genome/Gene_predictions and https://github.com/peterthorpe5/Heterodera_schachtii_genome_assembly. Genomic and RNAseq reads can be found under Bioproject: PRJNA722882.

## Author contributions

Flow cytometry: Sebastian Eves-van den Akker

Genome sample preparation and sequencing: Badou Mendy, Sebastian Eves-van den Akker

Data for assembly and/or gene prediction - Jose L. Lozano-Torres, Martijn Holterman, Joris J.M. van Steenbrugge, Mark G. Sterken, Johannes Helder, Tarek Hewezi, Tom R. Maier, Thomas J. Baum, Eric Grenier Assembly: Peter Thorpe, Sebastian Eves-van den Akker

RNAseq sample preparation and sequencing: M. Shamim Hasan, Clarissa Hiltl, Badou Mendy, Oliver Chitambo, Divykriti Chopra, Sebastian Eves-van den Akker

Differential expression and clustering: Clement Pellegrin, Helen Beasley, Olaf P. Kranse, Unnati

Sonowala Genome annotation: Peter Thorpe, Sebastian Eves-van den Akker

Effector predictions: Clement Pellegrin, Sebastian Eves-van den Akker

Horizontal Gene Transfer: Etienne G.J. Danchin, Georgios D. Koutsovoulos

Synteny/gene duplication: Rick E. Masonbrink, Peter Thorpe

Metabolic pathway analysis: Sebastian Eves-van den Akker

Mutant characterization and infection: Zoran S. Radakovic, Clarissa Hiltl, Esther Riemer

RNAi and qPCR: Zoran S. Radakovic, Samer S. Habash

*In situ* hybridisation: Samer S. Habash

Overexpression lines: Zoran S. Radakovic

ROS: Zoran S. Radakovic, Badou Mendy

Non-nematode infection assays: Zoran S. Radakovic

Plant phenotyping: Zoran S. Radakovic

Bacterial complementation: Zoran S. Radakovic, Nageena Zahid, Shahid Siddique, Julia Holbein

LM and TEM: Zoran S. Radakovic, Sławomir, Mirosław Sobczak

Ideas/conception: Shahid Siddique, Florian Grundler, Sebastian Eves-van den Akker,

Funding: Shahid Siddique, Florian Grundler, Sebastian Eves-van den Akker, Thomas J. Baum

Manuscript draft: Shahid Siddique, Sebastian Eves-van den Akker

Data management and supervision: Shahid Siddique, Florian Grundler, Sebastian Eves-van den Akker Comments: All authors

## Acknowledgments

The work at University of Bonn was supported by the Federal Ministry of Education and Research, Germany (BMBF) (Grant 031A326B to FMWG) and by the German Research Foundation (DFG) (Grant SI1739/3-1 and SI1739/5-1 to SS). MSH was supported by a fellowship from German Academic Exchange Service (DAAD). The work at University of California Davis was supported by the National Science Foundation (NSF) (Grant IOS-1954929) and National Institute of Food and Agriculture (NIFA) (Grant 20-3994). Work on plant-parasitic nematodes at the University of Cambridge is supported by DEFRA licence 125034/359149/3, and funded by BBSRC grants BB/R011311/1, BB/N021908/1, and BB/S006397/1. CP received funding from the European Union’s Horizon 2020 research and innovation programme under grant agreement No 882941. PT was supported by the University of St Andrews Bioinformatics Unit, funded by Wellcome Trust ISSF awards 105621/Z/14/Z and 204821/Z/16/Z. Work at Iowa State University was supported by Hatch and State of Iowa funds and a grant by the North Central Soybean Research Program. J.L.L-T was supported by a NWO domain Applied and Engineering Sciences VENI grant (14250) and a Wageningen University & Research Experimental Plant Sciences strategic funds grant. MGS was supported by NWO domain Applied and Engineering Sciences VENI grant (17282).

---

**Figure S1.**
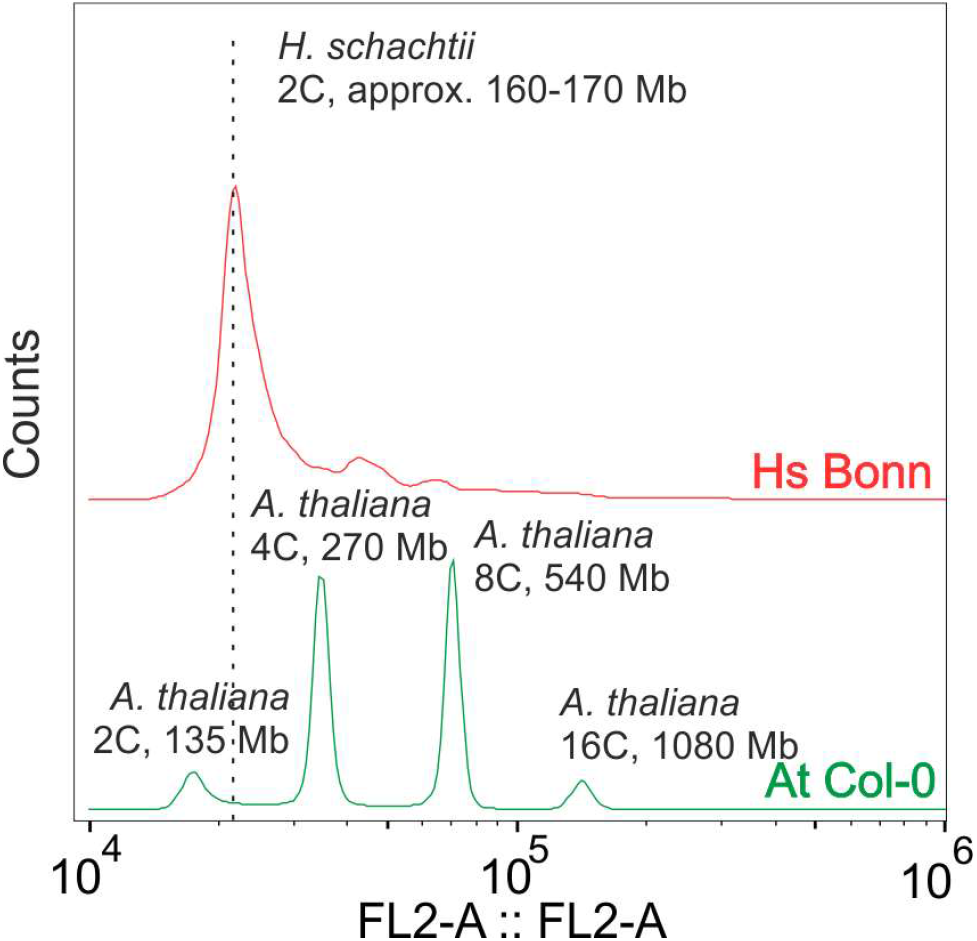
Flow cytometric estimation of genome size in *H. schachtii*. *A. thaliana* (Col-0 population) leaf (endoreduplication). Numbers in Mb represent genome length of collapsed haplotypes of each species. The *H. schachtii* (Hs Bonn population) data have been moved off the y-axis to allow ease of comparison.

**Figure S2.**
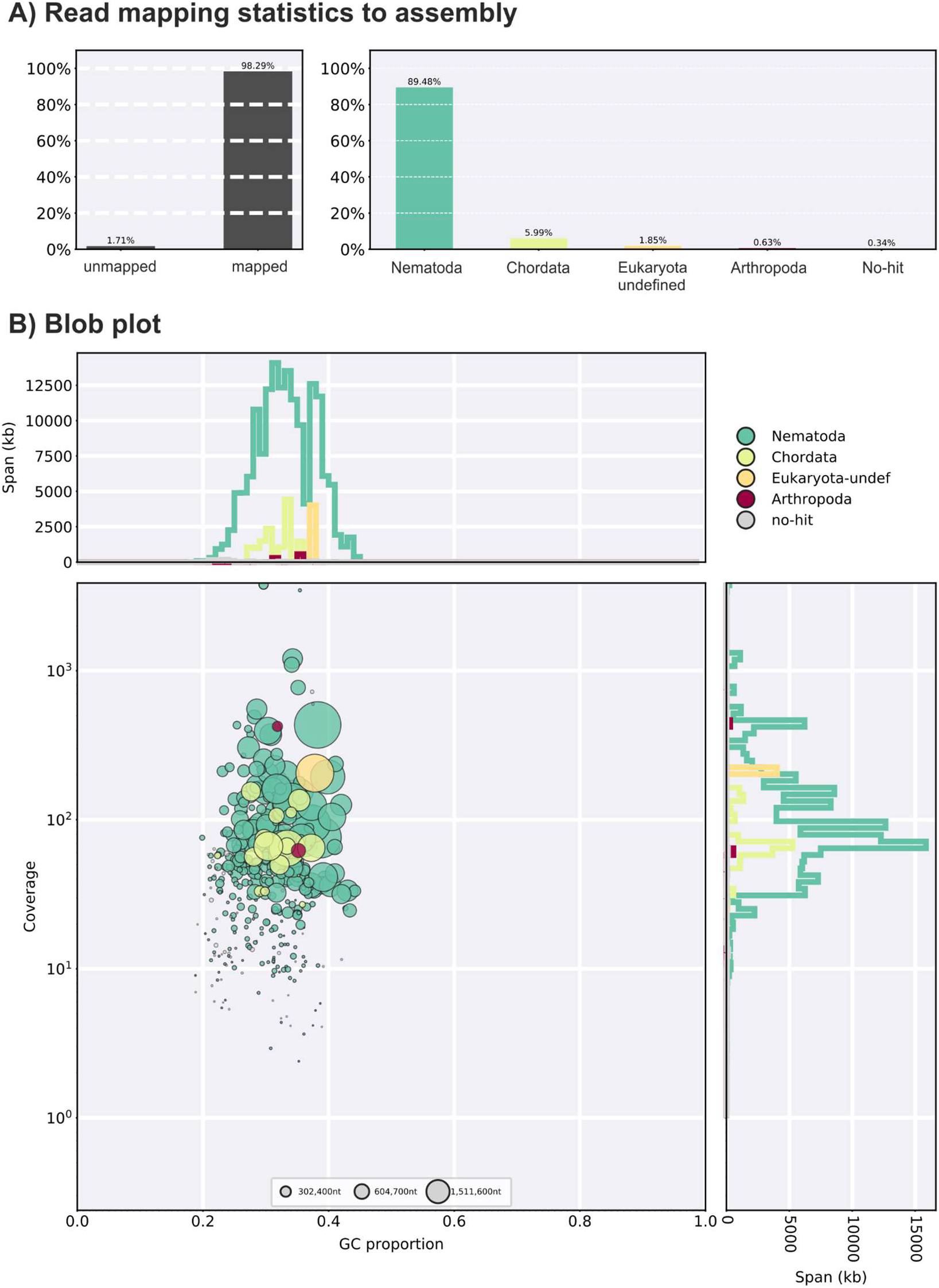
Blob plot contamination assessment of final assembly. **A**) Left, read mapping proportions to the final assembly of the *H. schachtii* genome. Right, distribution of mapped reads by taxonomic group. **B**) Twodimensional scatter plots, decorated with coverage and GC histograms. Circles are coloured by taxonomic group.

**Figure S3.**
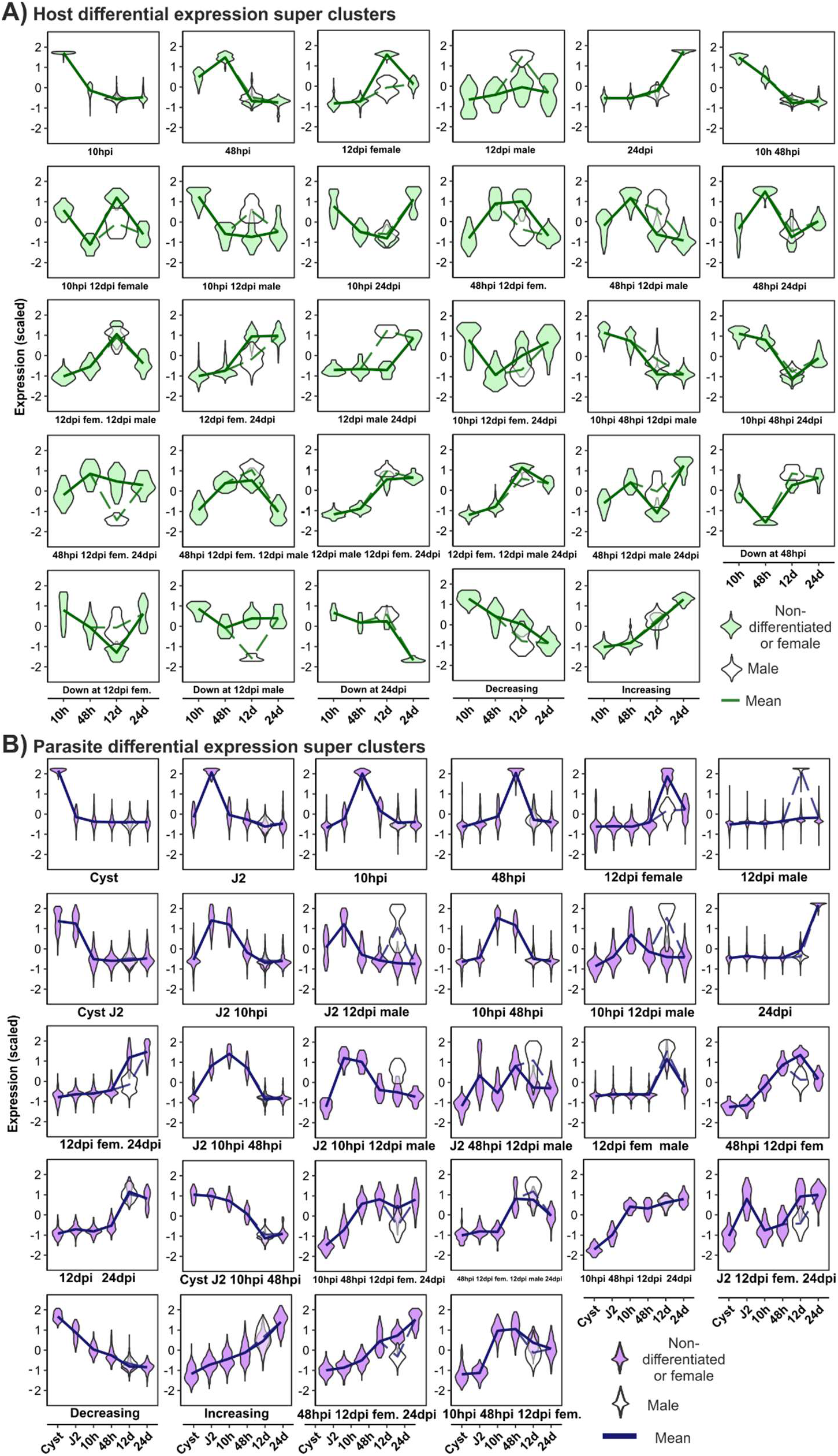
All differentially expressed super clusters. Expression super clusters for host (**A**) and parasite (**B**). Each violin subplot shows centred expression for genes in the cluster. The mean of each life stage is represented with a line. Open violins show gene expression for the 12 dpi male (or syncytial gene expression associated with it). Closed violins show gene expression of all other life stages (or the host gene expression associated with them).

**Figure S4.**
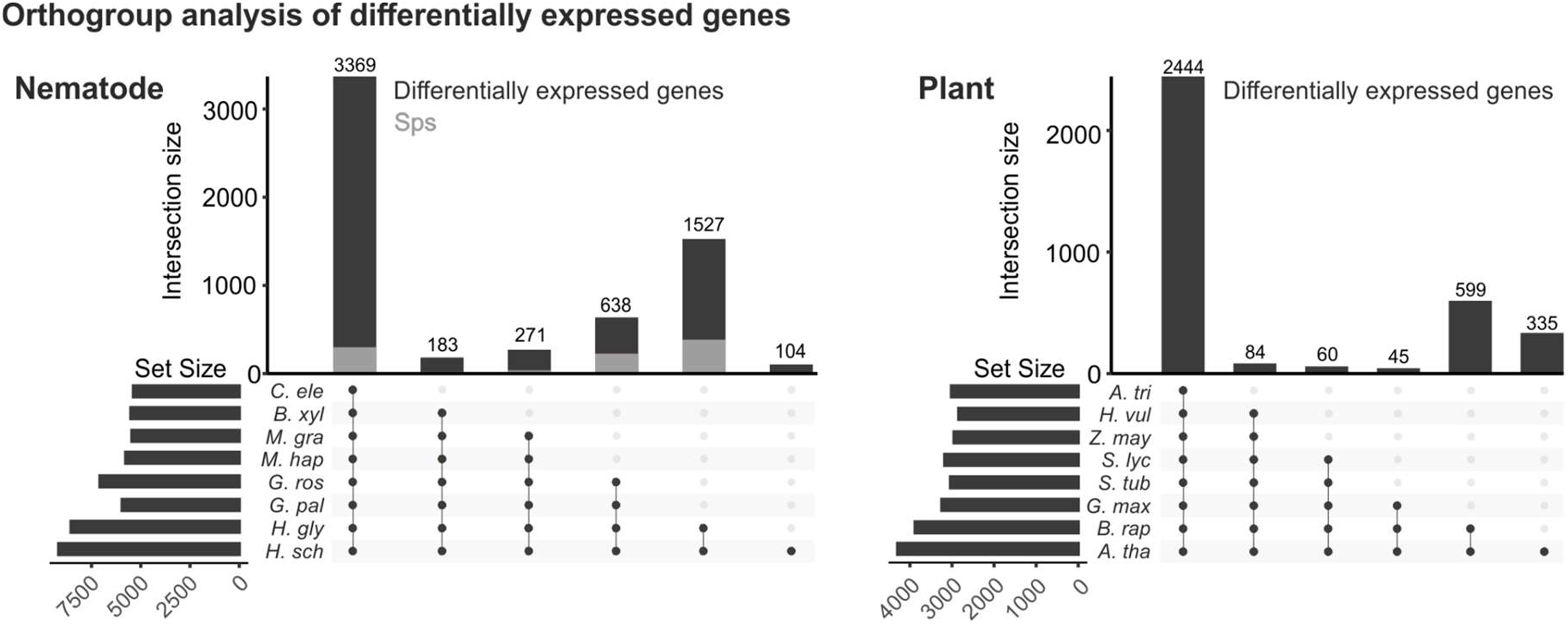
Orthogroup analysis of host and parasite. Upset graphs of orthologous gene clusters between eight nematode (left - *Caenorhabditis elegans, Bursaphelenchus xylophilus, Meloidogyne graminicola, M. hapla, Globodera pallida, G. rostochiensis, Heterodera glycines*, and *H. schachtii*) and eight plant species (right - *Amborella trichopoda, Hordeum vulgare, Zea mays, Solanum lycopersicum, S. tuberosum, Glycine max, Brassica rapa* (ssp. *rapa*) and *A. thaliana*). The intersections for 6 categories are shown. For example, there are 104 orthogroups that exclusively contain sequences from *H. schachtii*. Orthogroups that contain putatively secreted proteins (Sps) are indicated in grey.

**Figure S5.**
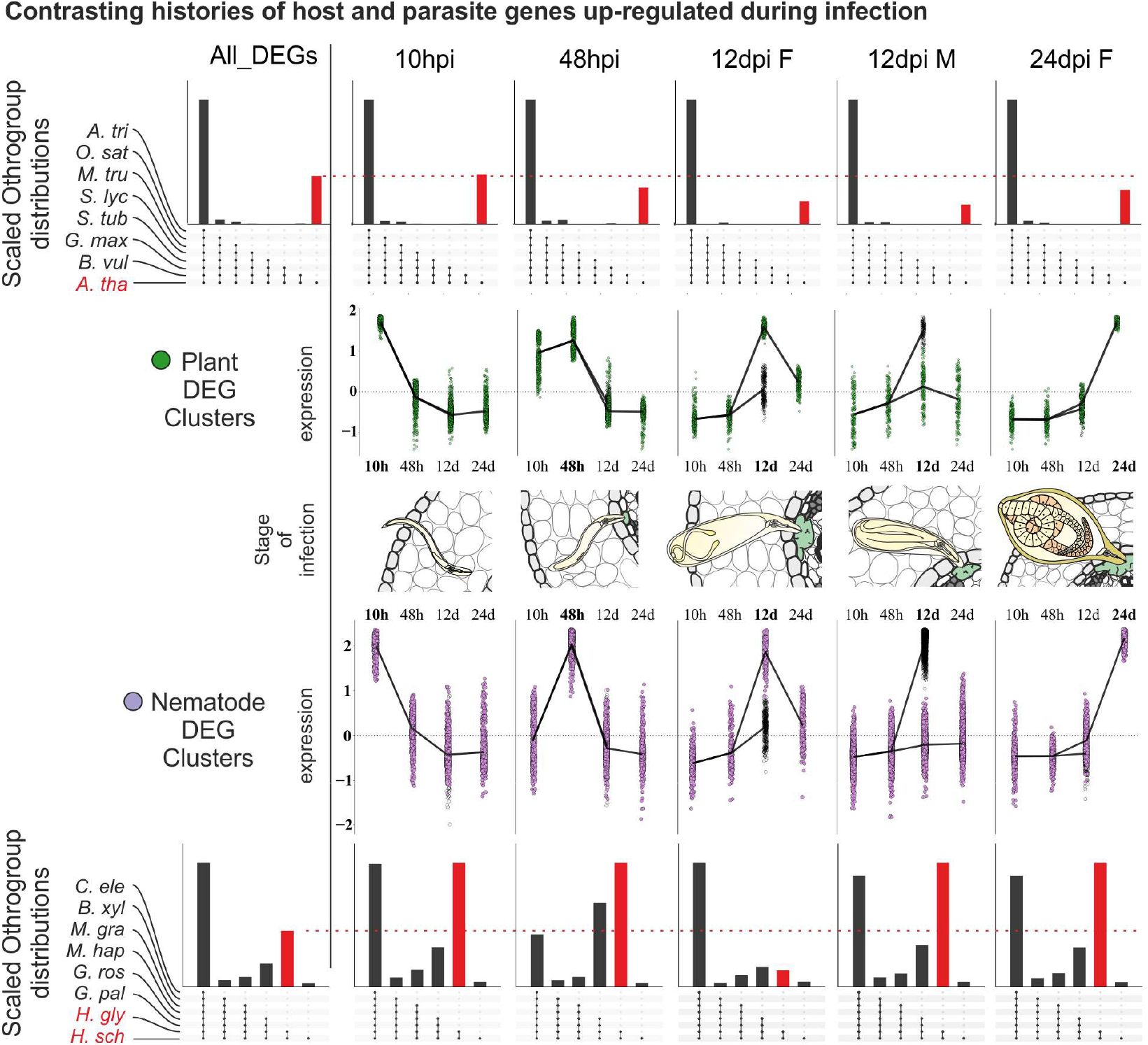
Contrasting evolutionary histories of host and parasite genes deployed at specific times of infection. Differential expression super clusters that describe discrete stages of infection (centre) for either the host (green) or the parasite (purple). Upset graphs of orthologous gene clusters between eight nematode (bottom - *Caenorhabditis elegans, Bursaphelenchus xylophilus, Meloidogyne graminicola, M. hapla, Globodera pallida, G. rostochiensis, Heterodera glycines*, and *H. schachtii*) and eight plant species (top - *Amborella trichopoda, Hordeum vulgare, Zea mays, Solanum lycopersicum, S. tuberosum, Glycine max, Brassica rapa* (ssp. *rapa*) and *A. thaliana*). Red highlights a subset of orthogroups for host and parasite.

**Figure S6.**
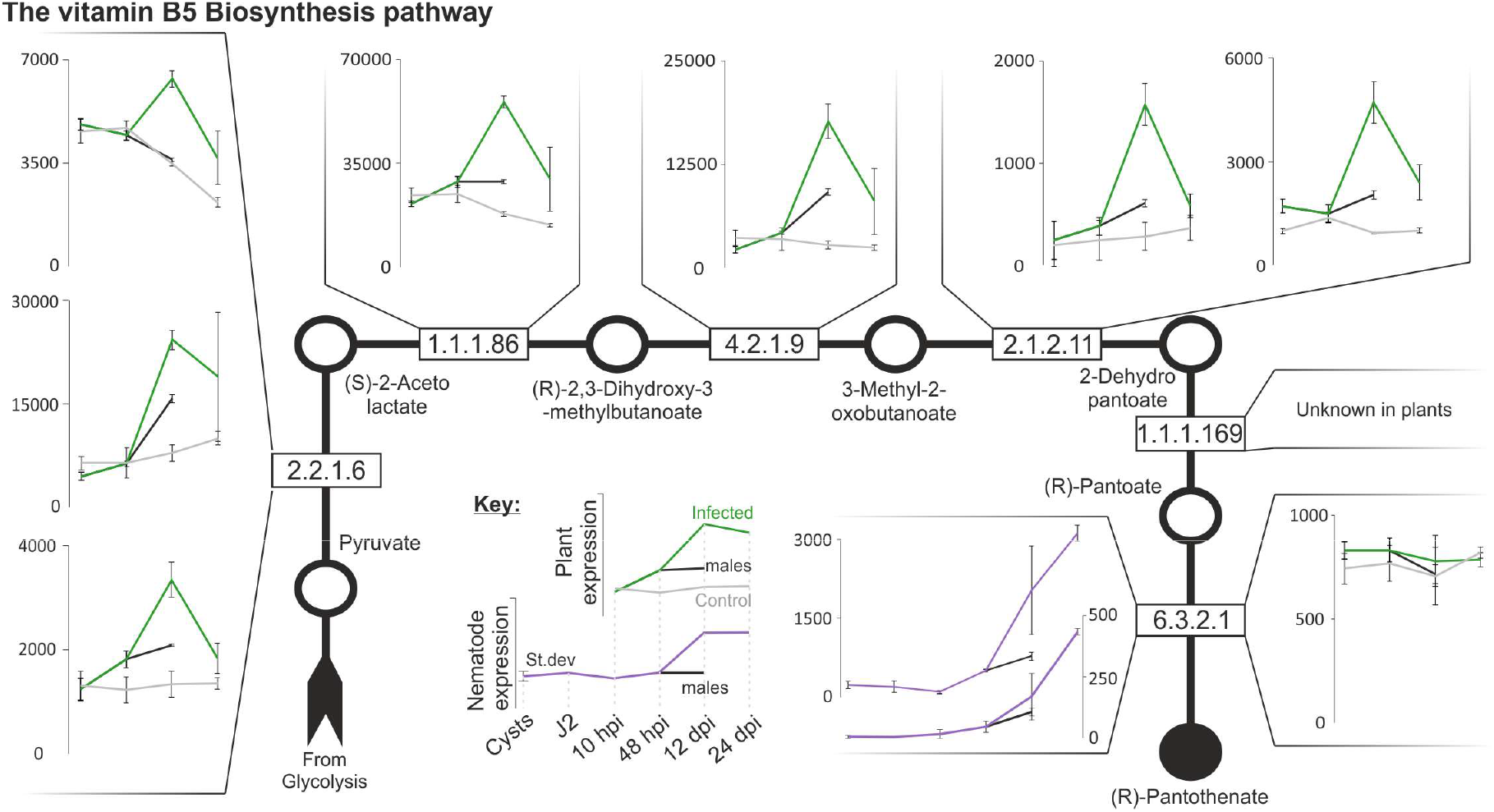
The complete vitamin B5 biosynthesis pathway. Black arrow indicates the linear pathway of vitamin B5 biosynthesis. Products/substrates are indicated with circles, enzymatic reactions (and corresponding EC codes) with squares. For each species, the expression profiles of the gene/s annotated with the EC codes at each step are shown (green for host, purple for parasite), if present. All steps are absent in the parasite *H. schachtii*, and present and upregulated at 14 dpi in the host *A. thaliana*, with the exception of the last step (PANC). PANC in the host is not upregulated at any time of infection, whereas PANC present in the parasite is upregulated during infection.

**Figure S7.**
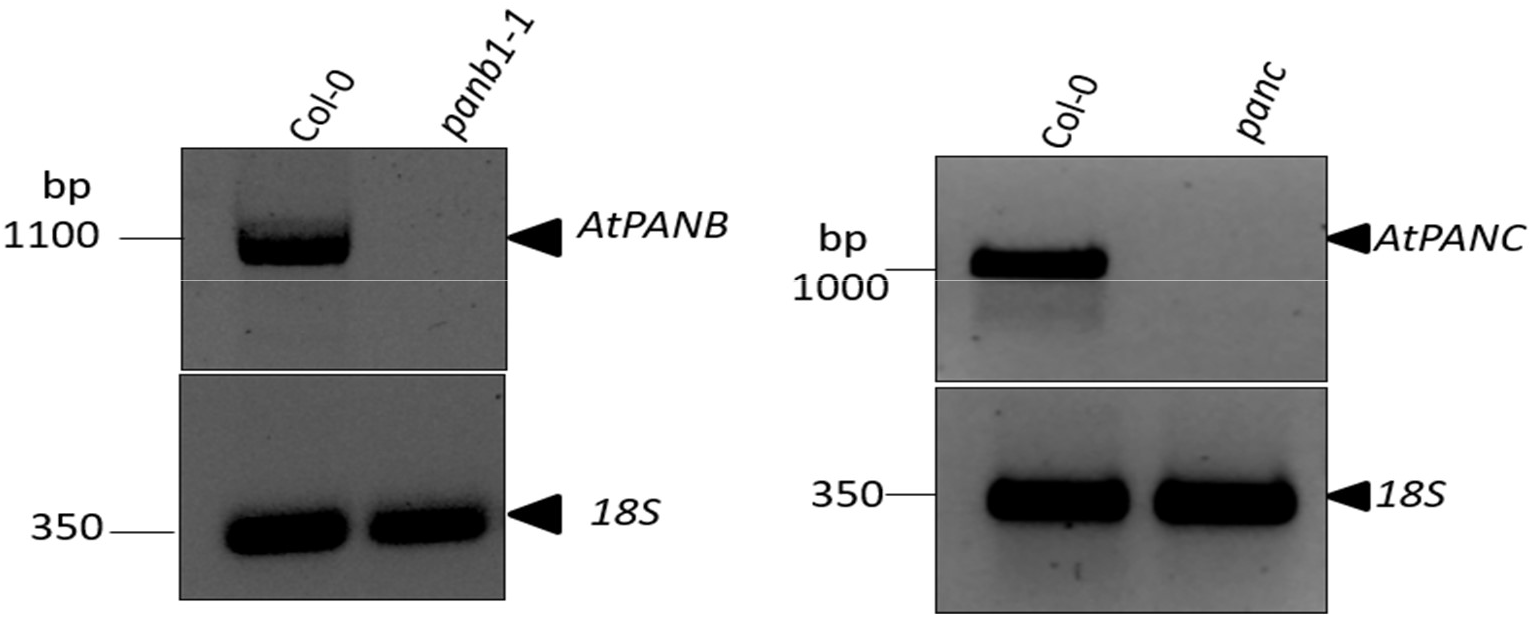
RT-PCR for presence or absence of *AtPANB* and *AtPANC* expression in wild-type or loss-of-functions mutant. RNA from Col-0, *atpanb1-1*, or *atpanc1* was extracted to synthesize single stranded cDNA. The presence or absence of expression is shown using primers given in Table S9. *18S* was used as a positive control.

**Figure S8.**
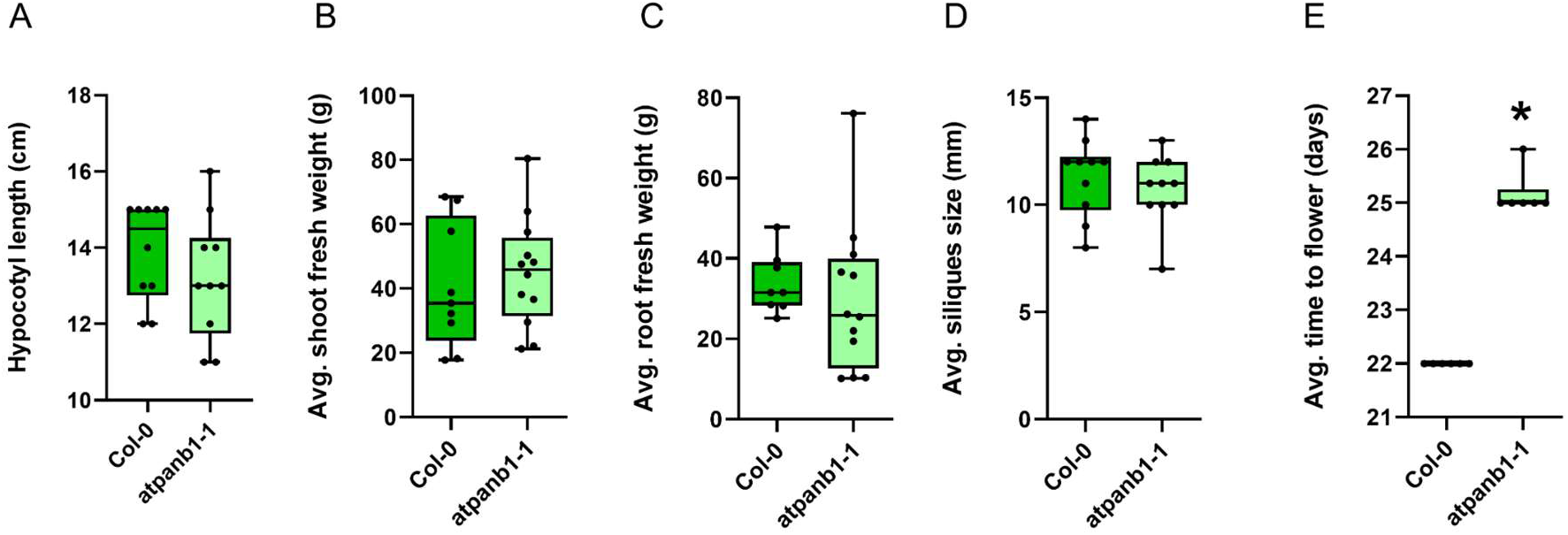
Phenotypic analysis of the Col-0 and *atpanb1-1* mutant. **A**) Average hypocotyl length of 5-days-old plants. **B**) Average shoot fresh weight of 12-days-old plants grown on Knop medium. **C**) Average root fresh weight of 12-days-old plants grown on Knop medium. **D**) Average silique length. **E**) Average number of days to flower. **D-E**) Plants were grown on Knop medium for 6 days before they were transferred to soil in the greenhouse.

**Figure S9.**
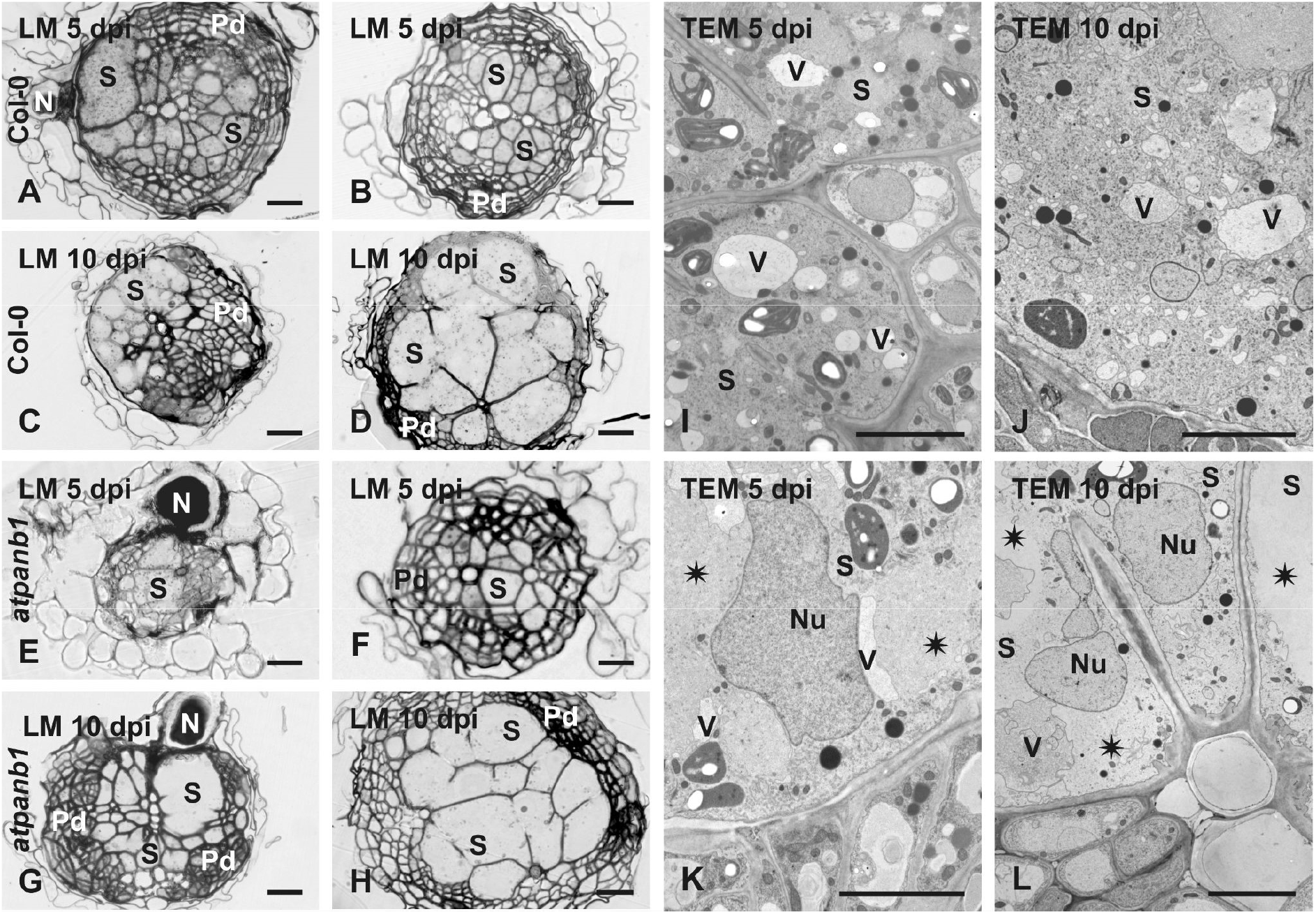
Development of syncytia induced by *H. schachtii* in Col-0 and *atpanb1-1* roots. Light (**A-H**) and transmission electron microscopy (**I-L**) images of sections taken from syncytia induced in roots of Col-0 (**A-D**, **I**, **J**) and *atpanb1-1* plants (**E-H**, **K**, **L**) grown on Knop medium without vitamin B5. Sections were taken from samples collected 5 (**A**, **B**, **E**, **F**, **I**, **K**) and 10 dpi (**C**, **D**, **G**, **H**, **J**, **L**). Light microscopy images (**A-H**) were made from sections taken in close vicinity of juvenile heads (**A**, **C**, **E**, **G**) or in some distance from the heads in the widest part of the syncytium (**B**, **D**, **F**, **H**). Transmission electron microscopy images (**I-L**) were obtained from sections taken at the widest part of syncytium. Asterisks indicate organelles-free regions of syncytial cytoplasm in *atpanb1* plants (**K**, **L**). Abbreviations: N, nematode; Nu, nucleus; Pd, periderm; S, syncytium; V, vacuole. Scale bars: 20 μm (**A-H**) and 5 μm (**I-L**).

**Figure S10.**
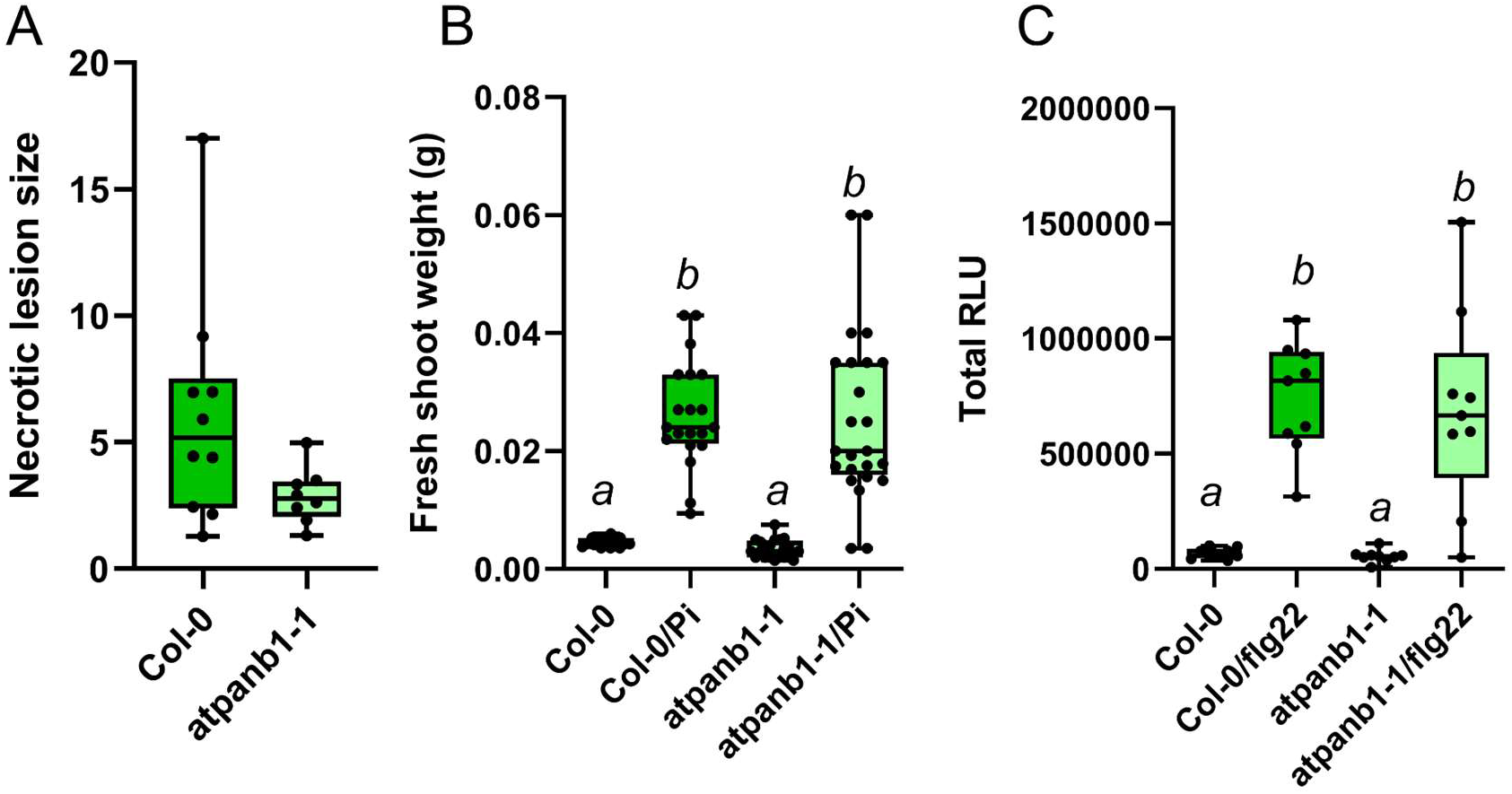
Loss-of-function *AtPANB1* is not impaired in plant immune responses. **A**) Infection assay with *Botrytis cinerea* under greenhouse conditions. Data were analysed using Student’s t-test. No significant difference was detected (95% confidence). **B**) Infection assay with *Piriformospora indica* for evaluation of its growth promoting effect. **C**) ROS burst in leaf disks treated with water or flg22. ROS burst was measured by using L-012 based assay from 0 to 120 min. **B-C**) Data were analysed using a one-way analysis of variance (ANOVA) followed by Tukey’s HSD post-hoc test. Bars represent mean ± SE. Columns not sharing the same letters indicate significantly different means (95% confidence).

